# Lack of functional STING modulates immunity but does not protect dopaminergic neurons in the alpha-synuclein preformed fibrils Parkinson’s Disease mouse model

**DOI:** 10.1101/2025.05.30.656977

**Authors:** Ida H. Klæstrup, Line S. Reinert, Sara A. Ferreira, Johanne Lauritsen, Gitte U. Toft, Hjalte Gram, Poul H. Jensen, Søren R. Paludan, Marina Romero-Ramos

**Author notes:** Corresponding author: Marina Romero-Ramos, Dept. of Biomedicine, Aarhus University, Hoegh-Guldbergsgade 10, DK-8000C Aarhus, Denmark.

## Abstract

Microglia response is proposed to be relevant in the neurogenerative process associated with alpha-synuclein (α-syn) pathology in Parkinson’s disease (PD). STING is a protein related to the immune sensing of DNA and autophagy, and it has been proposed to be involved in PD neurodegeneration. To investigate this, we injected 10 µg of murine pre-formed fibrils (PFFs) of α-syn (or monomeric and PBS as controls) into the striatum of wild-type (WT) and STING^gt/gt^ mice, which lack functional STING. We examined motor behavior and brain pathology at 1- and 6-months post-injection. STING^gt/gt^ mice showed more motor changes associated with PFF injection than WT mice. STING^gt/gt^ mice had a differential immune response to PFF with early and sustained increased microglia numbers and higher macrophagic CD68 response, but milder changes in the expression of immune-relevant markers such as TLR2, TLR4, IL1b, and TREM2. However, the lack of STING did not induce changes in the extent of α-syn pathology nor the p62 accumulation seen in the model. Altogether, this resulted in a faster but similar degree of nigrostriatal dopaminergic degeneration after 6 months. Therefore, the data do not support a necessary role for STING in the α-syn induced nigral neuronal loss in the PFF-PD mouse model used here. However, the results suggest a functional relevance for STING in the brain response to the excess and aggregation of amylogenic proteins such as α-syn that can contribute to symptomatic changes.

## Introduction

Parkinson’s disease (PD) is a neurodegenerative disorder characterized by significant loss of dopaminergic neurons in the substantia nigra (SN) and the presence of neuronal inclusions of aggregated alpha-synuclein (α-syn), Lewy bodies, through the central and peripheral nervous system (1). During PD, α-syn progressively transforms from a soluble protein to an insoluble amyloid, resulting in the characteristic pathology, which has been suggested can propagate to anatomically connected areas (2). This spreading seems to occur through the active release of modified α-syn and subsequent uptake by neighboring cells, where α-syn acts as a template and seeds aggregation-promoting pathology (3, 4). In parallel, α-syn-induced neurodegeneration comprises an inflammatory immune response. PD patients have an ongoing immune response that parallels, or even precedes, neurodegeneration (5). Patients exhibit elevated levels of pro-inflammatory cytokines and microgliosis related to α-syn pathology (6–9), which is suggested to play an active role in disease progression. α-syn is proposed to be a *danger-associated molecular pattern* initiating inflammation and promoting neurodegeneration (5). Aggregated α-syn can be recognized and bind to receptors on microglia, such as the toll-like receptors (TLR) 2 and TLR4 (10–14), thereby activating downstream pathways such as the NF-κB signaling and NLRP3 inflammasome assembly (10). Thus, α-syn plays a central role in the neuronal changes in PD and in the immune response that will contribute to disease progression. In addition, microglia are efficient in clearing out the released extracellular α-syn (15). However, the clearance efficiency depends on the microglia activation state and the α-syn solubility (15, 16). Therefore, the microglia also play a role in the α-syn spreading process.

Cyclic GMP-AMP synthase (cGAS) is a cytosolic DNA sensor that activates the innate immune system in response to double-stranded DNA in the cytoplasm, such as viral DNA (17, 18). The binding of DNA leads to the activation of Stimulator of interferon genes (STING), which induces the expression of type I interferons (IFNs) and NF-κB pathway genes, leading to antiviral responses (19–21). The cGAS-STING pathway can also be initiated by the presence of self-DNA, such as mitochondria and genomic DNA (17). Aggregated α-syn has been suggested to induce DNA damage, including double-strand breaks (22–24). However, it remains unclear whether this is sufficient to activate the cGAS-STING pathway. In addition, cGAS-STING activation during viral infection has been associated with a non-canonical autophagy activation (25, 26). While changes in autophagy have been proposed to play a central role in PD (27), α-syn pathology in models is consistently associated with the accumulation of the p62 (SQSTM1) adaptor protein (28).

In this study, we investigated the role of STING in the α-syn induced innate response and the associated neurodegenerative process. To do so, we injected murine α-syn pre-formed fibrils (PFF) into the striatum of wildtype (WT) and STING-deficient mice (STING^gt/gt^), or non-pathological monomeric (MONO) α-syn or PBS as controls. We examined the motor behavior, the central immune response, and the α-syn pathology at short-term (1 month) and long-term (6 months) post-injections (p.i). Our data suggest that STING contributes to the modulation of the immune response to both monomeric and fibrillar α-syn, suggesting that it is involved in the response to protein overload and α-syn aggregation. The lack of STING resulted in behavioral changes and accelerated the loss of dopaminergic axons. However, it did not lead to protection against the accumulation of pathological α-syn and neurodegeneration in the long-term.

## Results

### Lack of functional STING leads to different behavioral deficiencies

To explore the *in vivo* role of STING in the immune response and neurodegenerative process induced by α-syn pathology, MONO α-syn or α-syn PFFs were injected into the striatum of either wild-type (WT) or STING-deficient mice (STING^gt/gt^). As a control, PBS was injected into age-matched WT mice. To determine if the lack of STING influenced the behavioral phenotype in the PFF model, we examined the sensorimotor performance in two drug-free tests at 1- and 6-months p.i.

At 1 month p.i in the Challenging Beam test, all MONO-injected animals (WT and STING^gt/gt^) took significantly longer time to transverse the beam (Fig. 1A), made more errors (Fig. 1C), and took fewer steps/second (Sup. Fig. 1A) than PBS-treated mice, while MONO-WT also took more steps (vs PBS-WT) (Fig. 1B). Thus, the injection of monomeric α-syn, resulting to an excess of a soluble protein, also affected motor behavior. Regarding the PFF-injected group, only STING^gt/gt^ mice showed significant changes and took longer time (Fig. 1A) and fewer steps/second (Sup. Fig. 1A) than PBS-STING^gt/gt^. Furthermore, PFF-STING^gt/gt^ took more steps (Fig. 1B) and fewer steps/second (Sup. Fig. 1A) than PFF-WT, indicating a slower gait with smaller steps. Therefore, PFF injection particularly affects STING^gt/gt^ motorically at early stages.

**Figure 1.**
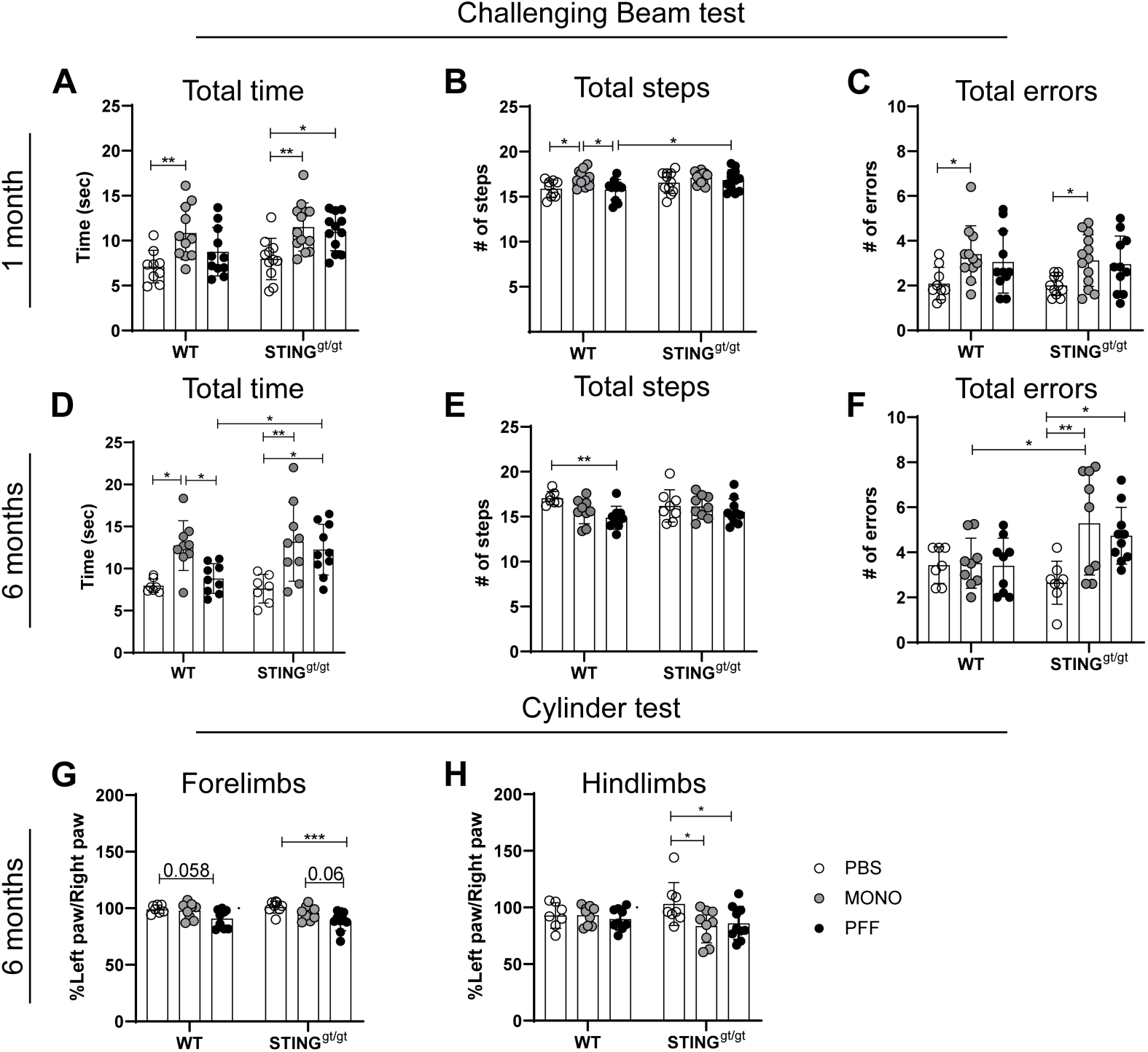
Motor behavior was assessed using the challenging beam and cylinder tests: Motor coordination and performance were evaluated by the challenging beam. **A-F)** Results from 1 month p.i; A) Total time to transverse the beam B) number of steps, and C) number of errors. Results from 6 months p.i; D) Total time to transverse the beam, E) number of steps, and f) number of errors. The figure displays the average results of frames 2, 3, and 4. Values are mean +SD. 1-month n=9-13, 6-months n=7-10. **G-H)**. To examine motor asymmetry cylinder test was evaluated 6 months post-injection, the percentage left paw use of the total right was calculated for the G) forelimbs and H) the hindlimbs. n=7-10. Two-way ANOVA followed by Sidak’s multiple comparisons. (See further stats in supplementary table 3). Data are mean +SD. *<p=0.05, **p≤0.01, ***p≤0.001, ****p≤0.0001.

After 6 months p.i, some of the behavioral changes were conserved. MONO-injected mice (WT and STING^gt/gt^) still took more time to transverse the beam (Fig. 1D) and walked fewer steps/second (Sup. Fig. 1B) than their respective PBS control. Notably, MONO-STING^gt/gt^ made more errors (Fig. 1F) and had more errors/step (vs. PBS) (Sup. Fig. 1B). In contrast, PFF-injections affected both genotypes differently; while PFF-WT did not significantly differ from the PBS-WT, except in fewer number of steps (Fig. 1E); PFF-STING^gt/gt^ took significantly more time to transverse, had more errors and errors/step, and fewer steps/second than PBS- STING^gt/gt^ (Sup. Fig. 1B). In fact, PFF-STING^gt/gt^ mice took longer (Fig. 1D) and, as in the 1- month time point, it had fewer steps/second than the PFF-WT mice (Sup. Fig. 1B). Equally, MONO-STING^gt/gt^ had worse motor coordination than MONO-WT, as it showed more errors (Fig. 1F) and errors/step (Sup. Fig. 1B).

The cylinder test at 6 months p.i showed a decrease of the activity in the PFF-WT mice, with fewer rearings and less use of the left (contralateral) forelimbs and hindlimbs (Sup. Fig. 2A, C and D), although only a trend towards asymmetry was seen in the forelimbs (p=0.058, vs. PBS-WT) (Fig. 1G&H). However, PFF-STING^gt/gt^ did not show differences in rearing, but showed a clear asymmetry using more touches with their right (ipsilateral) forelimbs and hindlimbs than the left (contra), which was confirmed by the lower percentage of the use of both left limbs (Fig. 1G&H and Sup. Fig. 2A). In conclusion, PFF injections induced greater motor changes in the STING^gt/gt^, indicating that a lack of STING signaling affects the brain response to α-syn.

### PFF injection leads to microgliosis and increased CD68 expression

To evaluate the microglia response, Iba1+ cells in SN were stereologically quantified, and morphological profiles were assigned in parallel as before (Fig. 2A) (29). Four profiles were defined: type A (resting or surveilling); no visible cytoplasm, small round dense nucleus, and long thin processes with little branching. Type B (hyper-ramified); visible cytoplasm, round dense nucleus, and many long-branched processes. Type C (hypertrophic); elongated and irregular cytoplasm, enlarged and less defined nucleus, shorter processes with varying thickness and less branching than type B. Type D (ameboid); big cell body merging with the processes, the nucleus fills most of the cells body, and the processes are very short, thick and few (30). While type A and B are most abundant microglia type in the brain, the type C constitute a small percentage while type D is very rarely seen in a healthy brain.

**Figure 2.**
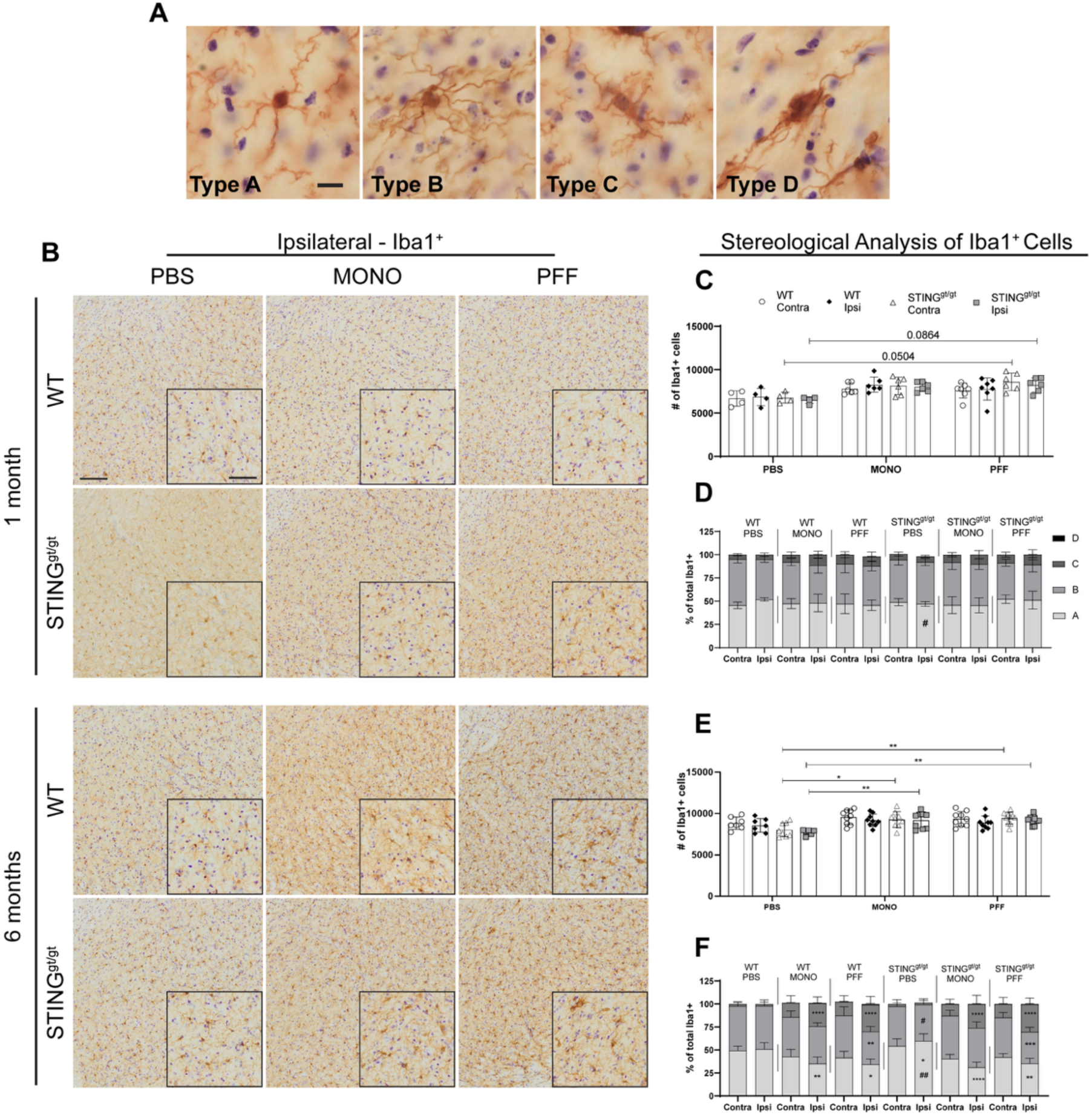
Microglia Iba1+ in the substantia nigra. **A)** Representative photos of the four Iba1+ morphological phenotypes quantified. Scalebar=10µm. **B)** Representative image of Iba1 staining in Substantia nigra (SN, both compacta and reticulata). Scalebar=100µm for all low magnification images, and 50µm for inserts. **C and E)** Stereological quantification of total Iba1 cells in the SN at 1-month (C) and 6 months post-injection (E). Three-way ANOVA with matched values followed by an uncorrected Fisher’s LSD with Bonferroni correction applied. Corrected values are displayed on the graphs (*<p=0.05, **p≤0.01, ***p≤0.001, ****p≤0.0001). **D and F)** The distribution of the morphological profiles are displayed as average percentage of total Iba1+ at 1 month (D) and 6 months post-injection (F). Two-way ANOVA (matched values when sides in the same group, or not matched values same sides and treatment) followed by Sidak’s multiple comparisons. * vs. the contralateral side within the same group and subtype. # vs. to the same side and treatment across genotypes. P #/*< 0.05, ##/**≤0.01, ###/ ***≤0.001, ####/****≤0.0001. (see further stats in supplementary table 3) Data are average ±SD, n=4-10.

At 1 month p.i, no significant difference was found in the total number of Iba1+ cells in the SN between all WT groups (Fig. 2B&C). However, we observed a bilateral increasing trend in Iba1+ cell number in PFF-STING^gt/gt^ (vs. PBS-STING^gt/gt^). Morphological distribution showed a decrease in the percentage of type A surveillant microglia in the STING^gt/gt^-PBS compared to WT-PBS (Fig. 2D (p=0.02)) with no other differences across groups (Sup. Fig. 3A).

In contrast, at 6 months p.i, we observed a significant bilateral increase in the number of Iba1+ cells in the STING^gt/gt^ mice injected with MONO or PFF (vs. PBS, Fig. 2B&E), while no differences were seen in the WT groups. Thus, α-syn injection, irrespective of the type/solubility, induced SN bilateral microgliosis in the STING^gt/gt^ mice but this tended to be faster in the PFF group.

Injections of α-syn led to changes in the distribution in microglia profiles in both genotypes after 6 months. The percentage of surveillant type A Iba1+ cells significantly decreased, while the percentage of hypertrophic Type C was increased in the ipsilateral SN (vs. contra) in both MONO and PFF-injected (Fig. 2F)). Interestingly, only PFF-injected mice, both WT and STING^gt/gt^, showed a significant decrease in the ipsilateral percentage of hyper-ramified type B cells (vs. contra) (Fig. 2F). As in the 1-month group, PBS intra-striatal injection also affected the STING^gt/gt^ microglia profile with increased type A and a decrease of type B cells in the ipsilateral side (Fig. 2F).

Our data -when expressed in total number of cells- showed that changes in microglia profile were also occurring in the contralateral SN in all α-syn injected animals, with more Type C cells irrespective of their genotype (Sup Fig 3B). In addition, while the ameboid Type D was never found in the PBS animals, and rarely in the α-syn injected mice at 1 month, it was more commonly seen in the α-syn mice after 6 months especially in the ipsilateral SN of PFF group irrespective of the genotype (Sup Fig. 3). Therefore, SN microglia, irrespective of the genotype and the α-syn solubility (MONO or PFF), responded to striatal α-syn injections by changing their profile towards a more hypertrophic and ameboid state. However, only the PFF injected mice showed a reduction of both the surveillant type A and hyper-ramified type B cells. Notably, in the STING^gt/gt^ mice, this was parallel to microglia proliferation, which was faster for PFF- than MONO-injected STING^gt/gt^ mice.

**Figure 3:**
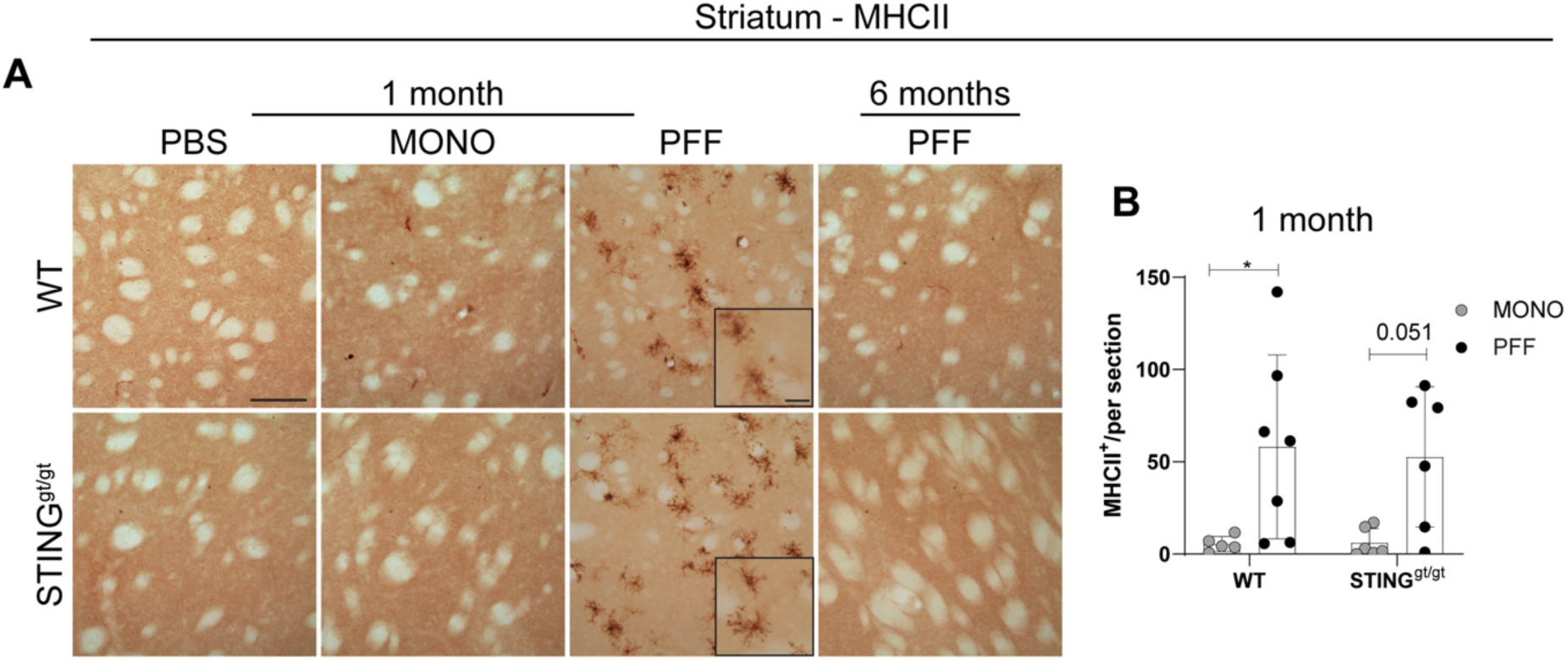
Analysis of immune marker MHCII. **A)** Representative images of MCHII staining in the striatum at 1- and 6-months post-injection. Scalebar=100µm, applies to all low magnification images and 25µm for all inserts. Graphs show **B)** average number of ramified MHCII cells in striatum 1-month post-injection. All analyses are performed on the ipsilateral hemisphere. Data average ±SD, n=4-10. Two-way ANOVA followed by Sidak’s multiple comparisons. (See further stats in supplementary table 3). P *<0.05, **≤0.01, ***≤0.001, ****p≤0.0001.

We also analyzed the expression of MHCII and CD68 in the striatum. Homeostatic microglia do not express MHCII, and only rare MHCII+ cells are found in healthy brain, normally in perivascular macrophages. At 1 month p.i, PFF injections resulted in increased MHCII striatal expression in WT, and a similar trend was observed in STING^gt/gt^ (Fig. 3A&B). This returned to the control level in all groups after 6 months (Fig. 3A). However, we did not observe any significant MHCII expression in the SN at any of the time points (Sup. Fig. 4).

CD68 expression, considered a marker of phagocytic activity in microglia, significantly increased at 1 month p.i. in the striatum of PFF-STING^gt/gt^ compared to MONO and PBS- STING^gt/gt^ but also to the PFF-WT (Fig 4A &B). However, at 6 months p.i, the striatal CD68 expression was equally increased in all PFF-injected animals of both genotypes (v.s. PBS and MONO). At this point MONO α-syn injection also increased striatal CD68 expression, although to a lower extent than PFF (vs. PBS, Fig. 4A &C). In contrast CD68 expression in SN at 1 month did not differ across groups, but after 6 months only PFF-STING^gt/gt^ showed significant upregulation compared to both PBS- and MONO- STING^gt/gt^ grousp (Fig. 4D, E & F). Overall, the data suggest that the PFF α-syn injections resulted in an early and sustained proliferation of microglia and a stronger phagocytic microglia response in the STING^gt/gt^.

**Figure 4:**
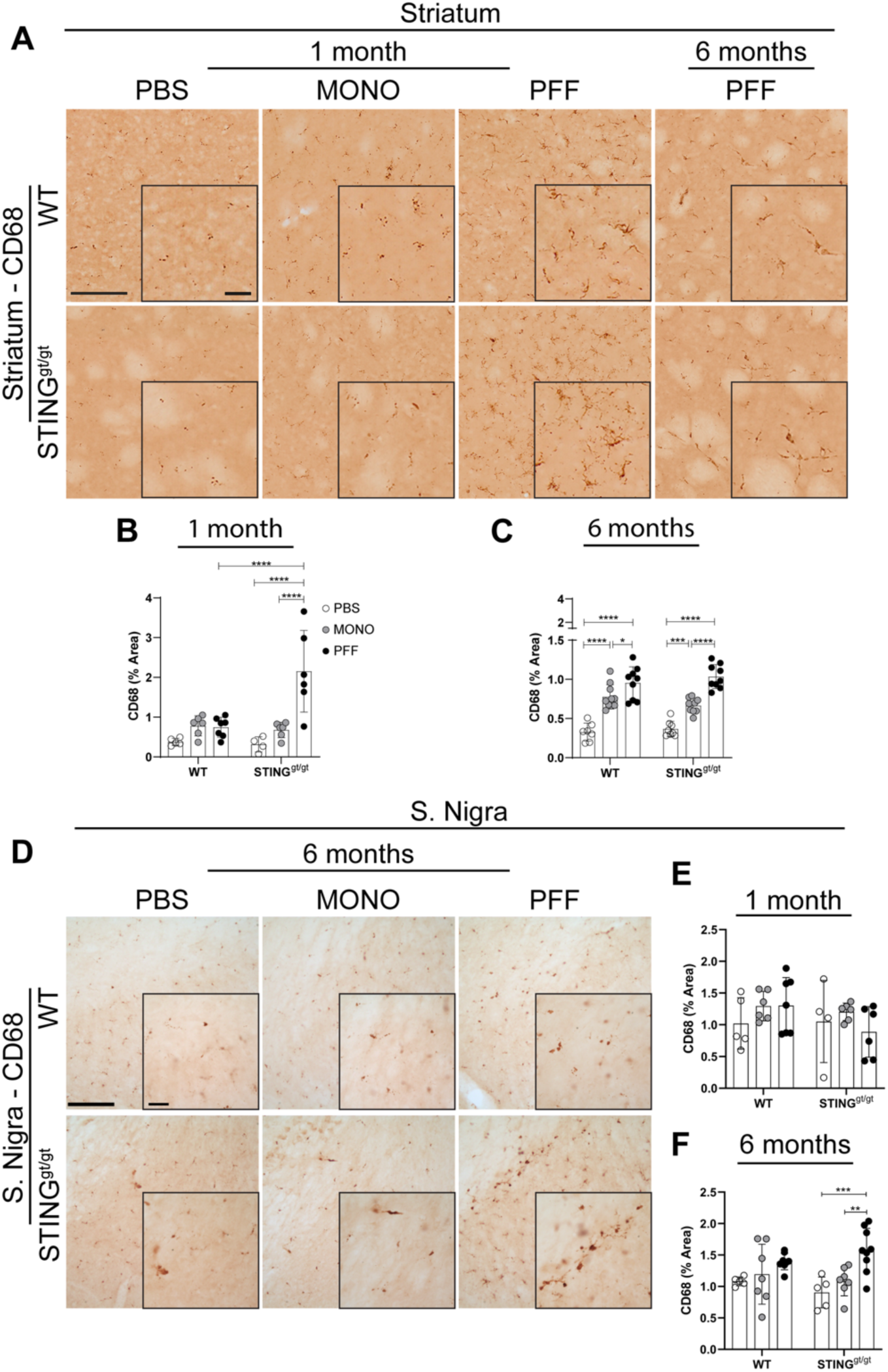
Analysis of immune marker CD68. **A)** Representative images of CD68 staining in the striatum at 1- and 6-months post-injection. Scalebar=100µm applies to all low-magnification images and 25µm for all inserts. **B and C)** Percentage of area covered by positive CD68 staining in the striatum at 1-month (B) and 6-months (C) post-injection. **D**) Representative images of CD68 staining in the substantia nigra at 6-months post-injection. Scale bars represent 100µm for all low-magnification images and 25 µm for all inserts. **E and F**) Graphs show the percentage of area covered by positive staining in the substantia nigra at 1-month (E) and 6-months (F) post-injection. All analyses are performed on the ipsilateral hemisphere. Data average ±SD, n = 4-9. Outliers were identified using Grubb’s test with alpha = 0.05. Two-way ANOVA followed by Sidak’s multiple comparisons. P **≤0.01, ***≤0.001.

### PFF treatment affects RNA expression and macrophage activation

To further characterize the early immune response in these mice, we examined the RNA expression in three different brain regions (pre-frontal cortex, striatum, and ventral midbrain) at 1 month p.i in MONO and PFF-injected mice in both genotypes. In the pre-frontal cortex from PFF-WT mice (Fig. 45A), we observe a significant increase of TLR 2 and 4, complement factor C1q, and TREM2 as well as a similar trend for the apoptotic-related protein p53- upregulated modulator of apoptosis (PUMA) (vs. MONO-WT). From these changes only the elevated C1q was also seen in PFF-STING^gt/gt^ group, which also showed unique upregulation of CXCL10 (p=0.055) (vs. MONO-STING^gt/gt^ control). Moreover, while PFF-STING^gt/gt^ showed lower TLR4 and TLR2 expression than PFF-WT mice, MONO-STING^gt/gt^ animals showed higher TLR4, but lower CCL2, IL6 (p=0.053) and the IFN induced gene Mx1 (p=0.052) than MONO-WT.

In the striatum of PFF-WT IFN- β (p=0.056), IL-1β, TLR2, C1q, C4b, TREM2 and PUMA were upregulated compared to MONO-WT (Fig. 5B). But in the PFF-STING^gt/gt^, only TLR2, C1q, and Mx1 were elevated vs. MONO-STING^gt/gt^. Moreover PFF-STING^gt/gt^ showed significantly lower CCL2, IL-1β, TLR2, and TREM2 than PFF-WT. While MONO-STING^gt/gt^ had lower CXCL10 and IL-6 expression than MONO-WT.

**Figure 5:**
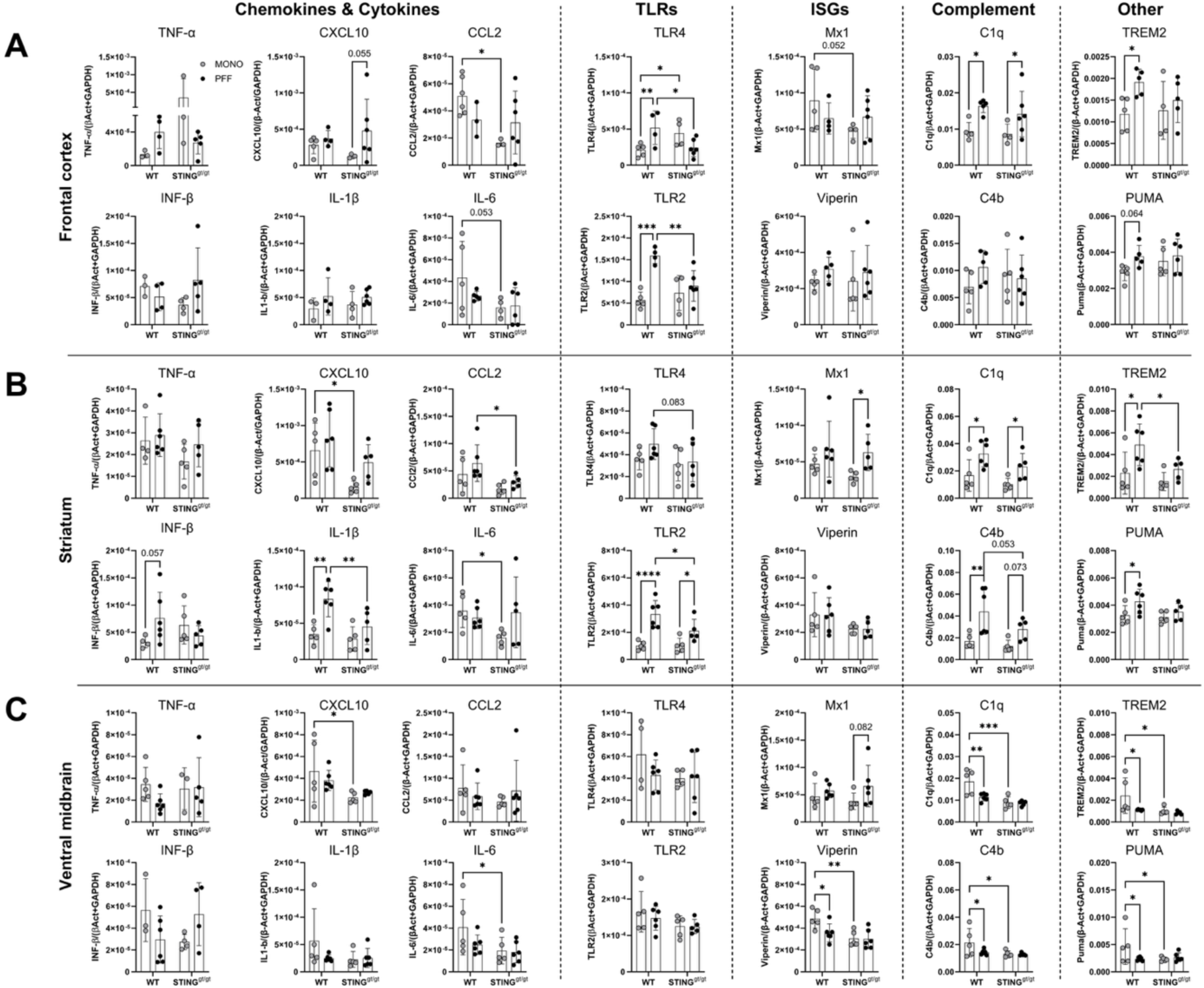
mRNA analysis at 1-month post-injection of PFF or MONO-in WT and STING^gt/gt.^ A-C) Results are displayed as the relative expression to the two housekeeping genes β-actin and GAPDH. For primers, see supplementary table 2. Two-way ANOVA followed by Sidak’s multiple comparisons. (See further stats in supplementary table 3). Data are average ± SD. n=4-6. *<p=0.05, **p≤0.01, ***p≤0.001, ****p≤0.0001

Conversely to the upregulation seen in the other two regions, the ventral midbrain of PFF-WT showed downregulation of Puma, C1q, C4b, Viperin, and TREM2 that seem to be a consequence of the higher expression of these transcripts in MONO-WT mice (Fig. 5C). Furthermore, we observed no difference between the two PFF groups (STING^gt/gt^ vs. WT), but a significant decrease in expression of CXCL10, IL6, C1q, C4b, Viperin, TREM2 and Puma in MONO-STING^gt/gt^ (vs. MONO-WT).

Together, the data indicate, first, that the early immune response to PFF intra-striatal injections is unique in the different brain regions, supporting an anatomically specific immune response. Secondly, the PFF-PD model leads to an early mRNA upregulation in the striatum and prefrontal cortex of well-known microglia activation-related markers involving TLR, complement system, and proinflammatory cytokines. Moreover, the lack of functional STING appears to reduce the cells’ ability to mount an immune response, both to the presence of monomeric and fibrillar α-syn.

### PFF injection leads to impaired autophagy and accumulation of pathological α-syn

To address whether STING might influence the PFF-induced α-syn pathology, we performed immunohistochemistry for aggregated (MJF14) and phosphorylated (p)Ser129 α-syn (Fig 6 & Supplementary Figs. 5&6). We observed immunostaining for both markers in regions anatomically connected to striatum as previously reported in the model such as: SN, amygdala, cortex and thalamic areas (Fig 6 and Sup. Fig 5&6). To study time progression, we selected two areas easy to delimit anatomically and with relevance in disease, namely: SN and amygdala. After 1 month, PFF injections resulted in significant aggregation of α-syn (MJF14+ staining) in the amygdala in WT (vs. MONO, supplementary Fig. 5 and Fig. 6B), with a similar trend in the STING^gt/gt^ (Fig. 6B). This aggregation was maintained after 6 months, when it became also significant in the SN for both genotypes (Fig. 6A-C). This was paralleled by accumulation of phosphorylated α-syn and 6 months after PFF injections pSer129 α-syn was significantly increased in both genotypes (vs. PBS and MONO) in the SN (Fig. 6G) frontal cortex and thalamus (Supplementary Fig. 6A-C), while in amygdala this was only significant in the PFF-STING^gt/gt^ but not in the PFF-WT (Fig 6F). However, no difference across genotypes was found suggesting that the lack of STING does not significantly modify the α-syn pathology nor its phosphorylation.

**Figure 6:**
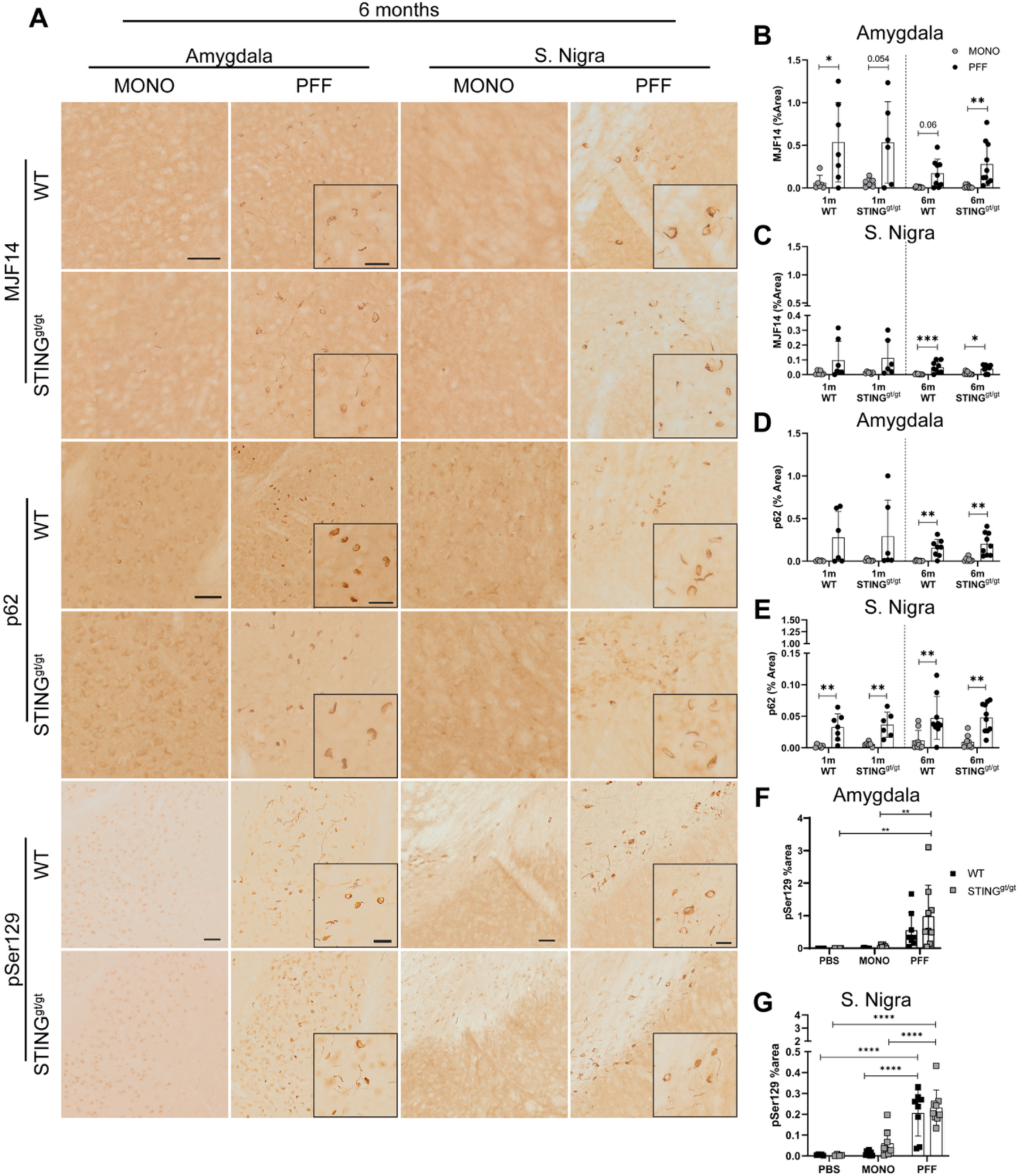
Analysis of pathology. **A)** Representative photos of MJF14, p62, and pSer129 staining in amygdala and substantia nigra 6 months post-injections. Scalebar=50µm for all low magnification in the same area and marker and 25µm for all inserts. **B, C)** Graphs show the percentage of area covered by MJF14 staining in the ipsilateral side at 1 and 6 months post-injection in the amygdala (B) and SN (C). **D, E)** Graphs show percentage of area covered by positive p62 staining at 1-month and 6-months post-injection in amygdala (D) and substantia nigra (SN). **F,G)** Graphs show percentage of area covered by pser129 staining in PBS, MONO, and PFF-injected animals 6 months post-injected in the amygdala (F), and SN (G) Two-way ANOVA followed by Sidak’s multiple comparisons (See further stats in supplementary table 3).). *<p=0.05, **p≤0.01, ***p≤0.001, ****p≤0.0001. All data are presented as mean ±SD. n= 6-10.

Since α-syn pathology has been associated with changes in autophagy, and because cGAS-STING can activate autophagy (25, 26), we measured the accumulation of the p62 adaptor protein (31). Parallel to our observations of α-syn pathology, at 1 month we found that p62 accumulated significantly in the SN but not the amygdala of PFF-treated mice regardless of genetic background (Fig. 6A, D, E & supplementary Fig 5). At 6-months p.i, p62 immunostaining was significantly increased in all PFF-treated mice (vs. MONO) in both the amygdala and SN (Fig. 6D and E). In conclusion, injection of PFFs induced pathological aggregation and phosphorylation of α-syn and p62 accumulation, which was not influenced by the lack of STING.

### Long-term PFF induced nigrostriatal degeneration is similar in WT and STING^gt/gt^ mice

To determine the integrity of the nigrostriatal pathway, we first evaluated the striatal dopaminergic terminals by densitometry analysis of tyrosine hydroxylase (TH) immunostained striatal sections. There was no statistically significant difference in the total number of TH neurons between the genotypes in the PBS groups (not shown). After 1 month, PFF injections induced a significant decrease in TH+ striatal signal in WT mice (vs. MONO) and STING^gt/gt^ (vs. MONO and PBS control) with a trend toward greater decrease than the one seen in the PFF-WT (p=0.07) (Fig. 7A and B). However, at 6 months p.i., the decrease of the TH+ striatal axonal signal was similar in the WT and STING^gt/gt^ in the PFF-injected groups, with a significant reduction compared to MONO and PBS for both genotypes (Fig. 7A and C).

**Figure 7.**
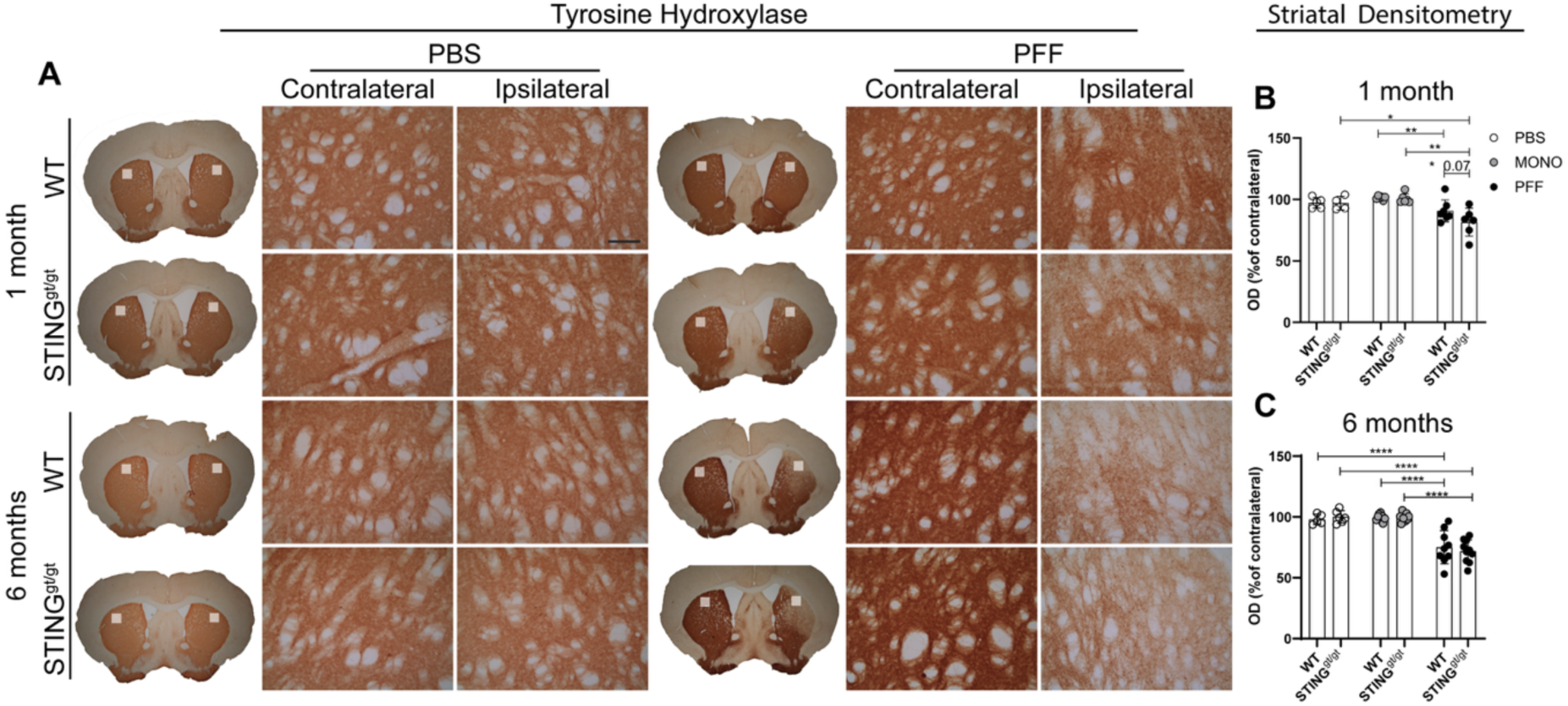
Loss of Nigro-striatal dopaminergic fibers in the striatum. **A)** Representative image of tyrosine hydroxylase (TH) immunostaining in the striatum 1- and 6 months post-injection. Scalebar=100µm apply to all. **B-C)** Densitometric analysis of TH+ in the ipsilateral striatal as the percentage of the contralateral at 1 month (B) and 6 months post-injection (C). Two-way ANOVA followed by Sidak’s multiple comparisons. Data are mean +SD. (See further stats in supplementary table 3). *<p=0.05, **p≤0.01, ***p≤0.001, ****p≤0.0001. SN=Substantia nigra.

Stereological quantification of nigral TH+ neurons at 1 month p.i showed no significant changes in WT mice (Fig. 8A). But, at that time point, both MONO and PFF-injected STING^gt/gt^ mice had a significant ipsilateral reduction of TH+ neurons in the SN compared to PBS (Fig. 8B). However, at 6 months p.i, both WT and STING^gt/gt^ PFF-injected animals had a similar TH+ neuronal loss (38-43%), which was significant to their respective MONO and PBS- controls (Fig. 8A and C). This data suggests an earlier dopaminergic degeneration in STING^gt/gt^, but a similar degree of neuronal loss at long-term, in both genotypes. It should be noted that since our analysis was based on TH expression, we cannot discard a possible downregulation of the enzyme, which constitute a limitation of our study.

**Figure 8.**
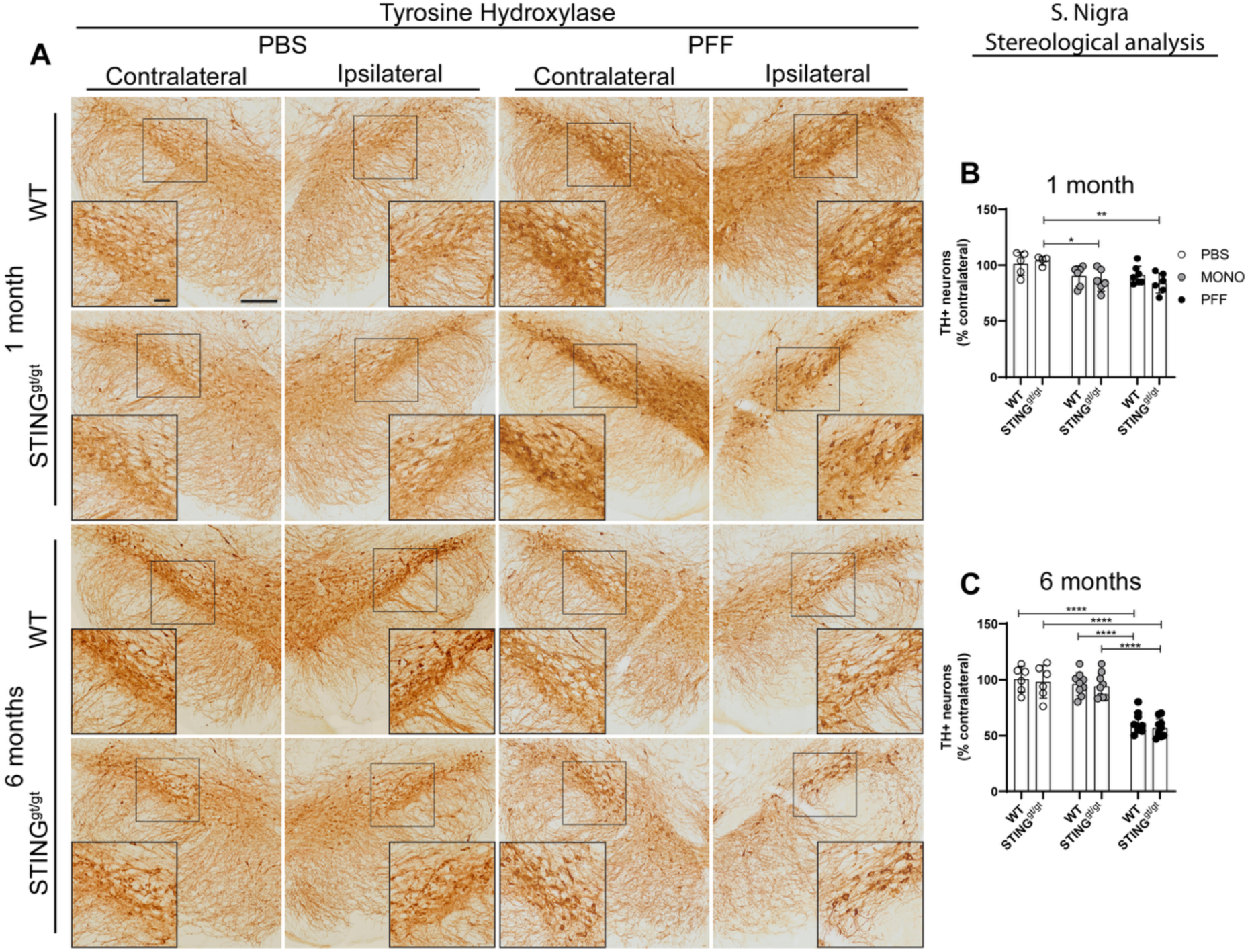
Assessment of tyrosine hydroxylase expressing neurons in substantia nigra. Lack of functional STING^gt/gt^ does not influence long-term degeneration of dopaminergic neurons**. A)** Representative image of tyrosine hydroxylase (TH) immunostaining in the SN at 1- and 6 months post-injection. Scalebar=200µm for low magnification images and50µm for inserts =**B-C)** Unbiased stereological quantification of the number of TH+ neurons in the ipsilateral SN as the percentage of contralateral at 1 month (n=4-10) (B) and 6 months (n=4-10) (C) post-injection. Two-way ANOVA followed by Sidak’s multiple comparisons (See further stats in supplementary table 3). Data are mean +SD. *<p=0.05, **p≤0.01, ***p≤0.001, ****p≤0.0001. SN=Substantia nigra.

## Discussion

STING is a signaling protein of the innate immune system involved in relaying signals from mislocated DNA to immunological effector functions, including inflammatory cytokine expression. STING is expressed in most cell types, including microglia in the brain (32). The activation of STING has been linked to two different mechanisms relevant to PD neurodegeneration: mitochondrial damage with subsequent release of mitochondrial (mit)DNA and the control of autophagic function (33, 34). For this reason, it has been proposed that STING might be involved in PD neurodegeneration. To investigate this, we injected 10 µg of murine PFFs α-syn (or MONO and PBS as controls) into the striatum of WT and STING^gt/gt^ mice, which lack functional STING. We examined motor behavior and brain pathology at 1- and 6-months p.i. (35). STING^gt/gt^ mice showed greater motor changes associated with PFF injection than WT mice, with longer time and more errors while transversing the challenging beam, and with asymmetry in the cylinder test. STING^gt/gt^ mice had a differential immune response to PFF than WT with early and sustained increases in microglia number and more robust macrophagic CD68 response, but milder changes in the expression of immune-relevant markers such as TLR2, TLR4, IL1b, and TREM2. However, the lack of STING did not induce changes in the extent of α-syn pathology nor the p62 accumulation seen in the model. Altogether, this resulted in a faster but similar degree of dopaminergic degeneration in the nigrostriatal system after 6 months. Therefore, our data do not support a necessary role for STING in the α-syn induced SN neuronal loss in the PFF-PD mice model used here. However, the results suggest a functional relevance for STING in the brain response to α-syn, both monomeric and fibrillar, which might result in changes in neuronal activity with meaningful phenotypic consequences in the motoric behavior.

Our data are in contrast with that previously published by Hinkle et al., who reported STING^gt/gt^ mice to be protected from the motor deficiencies, dopaminergic loss, and pathological α-syn accumulation induced by the PFF injections (36). When comparing both studies, the same mouse strains (C57BL/6 and STING^gt/gt^) and injection side (striatum) were used, and we both injected recombinant mouse α-syn PFF although different amounts per injection (10µg here vs. 2.5µg in theirs). To study progression and based in the literature we chose 1 month, to address early pathological and immune processes preceding neuronal loss, and 6 months as a timepoint with robust cell death and chronic neuroinflammation (37–39). However, in the Hinkel’s study 3 and 9 months p.i. were chosen. Our results here show that intrastriatal injections of α-syn PFFs induced Ser129 phosphorylation and aggregation of α-syn in multiple brain regions connected to the injection site: SN, amygdala, frontal cortex, and thalamus as before (35, 40, 41). But this was not influenced by the lack of STING in any of the areas analyzed. However, Hinkel’s study reported a lower percentage of pSer129 α-syn containing SN neurons in PFF-STING^gt/gt^ compared to PFF-WT (at 9 months p.i. with no other area nor any pathology marker analyzed). Since the induction of autophagy is a conserved function of STING (26, 42), serving as an antiviral defense and mediating p62-dependent attenuation of cGAS-STING signaling, we analyzed p62 (25). We found that the presence of pathological α- syn was accompanied by an accumulation of p62 in the same brain regions as seen before (28). This was not affected by the lack of functional STING, ruling out a key participation of STING-induced autophagy in the clearance of α-syn in our study. However, we observed a significant increase of CD68 expression in microglia in the striatum (1-month) and SN (6- months) of PFF-STING^gt/gt^. This might suggest a differential lysosomal activity in the absence of STING, as the glycoprotein CD68 is associated with the endosomal/lysosomal compartment (43). Thus, STING seems to be involved in the immune response to α-syn. This might be related to the earlier loss of dopaminergic SN neurons and dopaminergic terminals in the striatum in the PFF-STING^gt/gt^ that was significant already at 1-month p.i. However, the pathological aggregation of α-syn and long-term dopaminergic degeneration were similar across genotypes. Thus, our study does not support a crucial role for STING in the sustained SN neuronal death associated with α-syn aggregation, although it suggests a putative function for STING in the immune response to PFF-driven pathology.

PD and α-syn-induced neurodegeneration have been previously associated with inflammation mediated by TLR and pro-inflammatory cytokines (IL1β and TNFa). We measured RNA expression of a range of immune-related molecules in three different brain regions of PFF- bilaterally injected mice 1-month p.i. and used MONO as control. Our data confirmed the inflammation in the WT animals, with most of the transcriptomic changes found in the striatum (injection site) but also in the pre-frontal cortex, a distant anatomically related area that developed α-syn pathology. PFF-WT mice showed increased TLR2, TLR4, and IL1β, complement proteins C1q and C4b, and PUMA and TREM2, many of which have been previously related with the response of microglia during neurodegeneration and PD (44). This was damped -although not entirely avoided- in PFF- STING^gt/gt^ mice (vs. MONO), which also showed upregulation of TLR2 and C1q. This suggests a partial role for STING in the initial response to PFF, which is confirmed by the significantly lower IL1β, TLR2, and TLR4 expression in PFF-STING^gt/gt^ when compared to PFF-WT, indicative of a less inflammatory environment. Together, the data suggest that the STING does influence the early immune response to fibrillar α-syn. However, this was not sufficient to avoid nigral TH+ neuronal loss in our model. STING-induced inflammation is classically related to type 1 IFN signaling cascade. We did not observe robust significant changes in the PFF WT mice (vs. MONO) on IFN-related genes, such as Mx1 or Viperin, and IFNβ mRNA only showed a trend toward increase in the striatum. In contrast, Hinkel et al. showed increased mRNA of STING and other IFN-related genes in the striatum 3 months p.i. (vs. PBS as control), although they did not measure IFNs mRNA. We have previously reported that injections of MONO α-syn can induce an immune response without α-syn pathology nor neuronal death (45). Interestingly, we also observed differences between WT and STING^gt/gt^ in mice injected with MONO α-syn, with lower CCL2 (prefrontal cortex), CXCL10 (striatum & midbrain), IL6 (striatum & prefrontal cortex), PUMA, C1q, C4b, Viperin, and TREM2 (midbrain). Some of these proteins have been previously related to STING activation, such as Viperin, CXCL10, PUMA (46, 47), and TREM2, which we recently showed to be involved in the antiviral defense mediated by cGAS-STING in the brain (48). Notably, it has been suggested that TREM2 expression might exert certain protection toward Alzheimer-related neurodegeneration. Thus, this downregulation might be deleterious (49). All of this suggests a role for STING in the response to both monomeric and pathological aggregated α-syn, or potentially towards excess levels of proteins (as monomeric α-syn). This transcriptional response was parallel to a distinctive microglia profile changes towards more hypertrophic subtypes in the SN, suggesting certain long-lasting immune responses in both WT and STING^gt/gt^. However, we also observed microglial proliferation in the STING^gt/gt^, already at 1 month in the PFF and after 6 months in both MONO and PFF STING^gt/gt^ (vs. PBS) but not in the WT mice, further corroborating a differential microglial response. This, together with the increased CD68 expression in the microglia in PFF- STING^gt/gt^, suggests that the lack of STING influences the short- and long-term immune response associated with the PFF-PD model, by decreasing the expression of important pro-inflammatory proteins and by promoting early and sustained microgliosis and phagocytosis. These changes paralleled an earlier nigro-striatal degeneration but were not sufficient to modify the long-term dopaminergic SN loss.

STING hyperactivation leads to dopaminergic neuronal death, confirming that STING-related inflammatory stress is sufficient (50). Several articles provide a link between α-syn and IFNs, and increased α-syn levels have been associated with viral infections (51–53). α-Syn seems involved in type 1 IFN response against an RNA virus as it is increased during infection, and if α-syn was absent, the response failed (54). Accordingly, SARS-CoV-2 induced STING activation has been associated with α-syn upregulation (55). Thus, under certain conditions, repeated viral infection might increase the risk of developing α-syn pathology; accordingly, α- syn pathology has been seen in a rodent model of SARS-CoV-2 infection (56). Therefore, STING activation might be sufficient, but not necessary, for PD-like neurodegeneration, which might be mediated by inflammation and α-syn aggregation. On the other hand, IFNβ receptor signaling has been related to autophagy in neurons, and its absence led to α-syn neuronal accumulation (57), thus supporting a protective role for IFN signal. We did not see significant IFNβ changes nor differences in p62 accumulation, suggesting that this mechanism is not particularly relevant in the model used here. However, our analysis cannot discard possible changes in cellular machinery with functional consequences in neurons. Moreover, the differential immune environment in the STING^gt/gt^ will also influence neuronal activity. Indeed, it has been shown that microglia regulate neuronal activity with meaningful behavioral outcomes (58). Furthermore, both inflammatory and anti-inflammatory cytokines can modulate neuronal excitability and behavior (59), and chemokines can modulate synaptic plasticity and activity (60). Accordingly, we observed more motor alterations in the PFF-STING^gt/gt^ than PFF- WT, suggesting greater affection of the basal ganglia circuitry in the mice lacking STING. Interestingly, for PFF-WT, we do not observe a worsening in the challenging beam test performance with time, despite the significant loss of dopaminergic neurons at 6 months. This further corroborates that the behavioral deficiencies in the model are not solely indicative of neuronal death, but they can be at least partially associated with microglia/immune changes that in turn affect neuronal activity. This was corroborated by the motor defects in MONO-mice that were slower when transversing the challenging beam. Again, MONO-STING^gt/gt^ mice were more affected than MONO-WT, which we could speculate was related to the nigral microgliosis and phagocytic activity observed in this group. In conclusion, we observed more compromised motor capacity in STING^gt/gt^ mice despite having a similar degree of α-syn pathology and dopaminergic cell loss to WT. Therefore, the lack of STING resulted in differential response to monomeric and fibrillar α-syn with early and long-term microglia changes that had phenotypical motoric consequences.

The cGAS-STING pathway is involved in the response to the presence of viral or mitochondrial DNA in the cytosol. Age-associated failure in mitophagy has been shown to induce type 1 IFN activation, cGAS-STING inflammation, and the presence of mtDNA in cytosol (61). The relevance of mitochondrial dysfunction in PD is well accepted. However, the possible presence of DNA in the cytosol in sporadic (s)PD is not well studied. Due to their described function, Pink1 and Parkin-related PD are associated with mitochondrial damage (62). Increased circulating cell-free mtDNA was found in the sera of biallelic and heterozygous PRKN/PINK1 PD patients who also presented increased IL6, suggesting an association with inflammation (63). LRRK2(G2019S)-PD patients also showed elevated cell-free mtDNA in CSF vs. non- manifested carriers and sPD patients (64). However, no increase of cell-free mtDNA was seen in sPD in both studies (63, 64). Some in vitro studies suggest that the release of mtDNA might occur in other genetic PD forms. For example, the accumulation of glucosylceramides (relevant in GBA-PD) in microglia in vitro was associated with STING activation and mtDNA in the cytosol (65). Also, in vitro, mutated α-syn interaction with TOM40 leads to mitochondrial damage and mtDNA in cytosol (66). DNA damage response in neurons has been described in the AAV-human α-syn mice model and a (human) PFF-based model, which in vitro was associated with oxidative stress and mitochondrial dysfunction (36, 67). However, none of the studies investigated the presence of cytosolic mtDNA. Thus, despite the signs of DNA damage in cells with α-syn pathology and displaced mtDNA in genetic forms of PD, no definitive evidence of mtDNA in the cytosol has been shown so far in sPD. Even though we did not measure this, which is a limitation in our study, our data does not suggest the presence of cytosolic DNA in amounts sufficient to lead to STING-mediated inflammation that might be significant for neuronal death.

Our data contrasts with that from Hinkle et al.,(36) however, we should also note some differences in the production protocol for the recombinant mouse α-syn PFFs. Our PFFs consist of only the isolated insoluble material (pellet) post-aggregation vs. theirs, which includes both the soluble and insoluble material in their final “PFF stock”, which they state may contain 50% of non-aggregated α-syn (68). This may result in a smaller final dose of aggregated α-syn in their preparation vs. ours. There are also a few differences in purification (SEC & IEX vs. IEX & RP-HPLC) and storing (in solution at 30mg/mL vs. lyophilized) of the α- syn monomer used to produce the PFF, which may lead to residual impurities or oligomers that could affect batch-to-batch consistency in aggregation during PFF preparation. In addition, Hinkel et al. injected bilaterally 2.5µg (total 5 µg), whereas we performed unilateral injections of 10µg α-syn PFF, and they analysed behavior and dopaminergic survival at 9 months p.i while we did so at 1- and 6-months p.i. It is unclear the relevance of these collective differences in the model (69).

In conclusion, our study does not support a crucial role for STING in long-term α-syn pathological aggregation and the dopaminergic neuronal death in the murine-PFF-PD mice model. However, STING seems to be involved in the microglia response to both monomeric and fibrillar α-syn. The absence of STING lead to a differential immune environment in the PFF model, with decreased expression of pro-inflammatory mediators (TLR2 and 4 and IL1B), increased CD68-lysosomal expression, and early and long-lasting microgliosis. The differential microglia/immune status resulted in neuronal functional changes that have a relevant phenotypic/behavioral consequence, with STING^gt/gt^ mice exhibiting more motor defects. Therefore, STING seems to have a relevant function in the microglia response to the excess and aggregation of amylogenic proteins such as α-syn that can contribute to symptomatic change.

## Materials and Methods

### Animals

Male and female wildtype (WT) C57bL/6J mice (male=31, female=27), C57BL/6- Tmem173 <gt> “Goldenticket” (STING^gt/gt^) animals (male=30, female=31) were bread at Taconic. Animals were housed in a maximum of 4 per cage, with ad libitum access to water and food, and in a climate-controlled animal facility under a 12h/12h night/day cycle. At the time of surgery, animals were on average 2 months old and weighed 20-30g. All animal experiments were conducted under humane conditions, following the rules set by the research ethical committee at Aarhus University, and in is accordance with the guidelines established by the Danish Animal Inspectorate and Danish law.

### Protein purification and aggregation of mouse α-synuclein

Mouse α-syn were generated in our collaborator Dr. PH Jensen’s lab. The protein was recombinantly expressed in E. coli BL21 DE3 bacteria, using the pRK172 plasmid encoding for mouse ASYN (kind gift from Prof. Virginia Lee (University of Pennsylvania)). The bacteria were pelleted by centrifugation and resuspended in buffer (50 mM Tris, 1 mM EDTA, 0.1 mM DTE, 0.1mM PMSF, pH 7.0). The suspension was lysed by sonication in Branson Sonifier (15min, 70% power, 50ms on/off) and centrifuged at 20,000 g for 30 minutes at 4°C. The supernatant was boiled for 5 min and centrifuged at 48.000 g for 30 min at 4° C. Before ion exchange purification, the supernatant was dialyzed in 20 mM tris pH 6.5 and filtered through a 45µm filter. Mouse ASYN was purified on POROS HQ 50 ion exchange chromatography with a continuous gradient of 0% - 100% 2 M NaCl in 20 mM Tris pH 6.5. The fractions with mouse ASYN were isolated and further purified by reverse phase chromatography (C18) to remove nucleotides and lipids bound to ASYN, this step also removes endotoxins (lipoglycans) from the sample (<0.5 EU/mg, confirmed using the Pierce^TM^ Chromogenic Endotoxin Quant Kit, ThermoScientific^TM^). The pure protein was dialyzed in 20 mM ammonium bicarbonate, lyophilized, and stored at -20° C. To produce sterile preformed fibrils (PFF), mouse ASYN was solubilized in PBS (Gibco)) and passed through a 100kDa filter (Amicon Ultra) to remove unwarranted oligomer species, before sterile filtering through a 0.22um filter (Merck). Protein concentration adjusted to 1mg/mL and seeded with 5% (m/m) sonicated mouse alpha- synuclein pre-formed fibrils (produced previously and used before (70)). The solution was allowed to aggregate at 37° C, 1000 RPM for 2 days. The insoluble ASYN PFF were isolated from unbound monomeric ASYN by 15,600g centrifugation for 30 min at 20°C and resuspended in fresh sterile PBS (Gibco). The protein concentration was determined by the Bicinchoninic acid (BCA) protein concentration assay (Pierce). Aliquots were frozen at -80°C until the day of use, when PFF was sonicated for 40 min (0.5s off/0.5s on, 30% output power). To confirm correct sonication and fibril size, dynamic light scattering (DLS) was used (222- DPN, NanoStar). The optimal size range is between 20-50 nm (71). These DLS measurements were done before and after surgery to confirm the stability of the fibrils and maintenance of the optimal size.

### Animal surgery

WT, STING^gt/gt^, and cGAS-KO animals were both randomly divided into receiving either mouse monomeric α-syn, α-syn pre-formed fibrils (PFF) or Dulbecco’s Phosphate Buffered Saline (PBS) (Biowest). Mice were anesthetized with a combination of medetomidine hydrochloride (1mg/ml), midazolam (5mg/ml), and fentanyl (50µg/ml) in 0.9% NaCl solution in a final dose of 10ml/kg (0,4-0,5 ml/injection, i.p.) and placed in a stereotaxic apparatus. Intracranial injection of α-syn fibrils or monomers (2µl of total 10µg, diluted in PBS) or PBS (2µL) were performed. Injections were performed with a 5µL Hamilton syringe with a pre-made glass capillary attached. Unilateral animals were injected into the right hemisphere at the coordinates: A.P +0.7mm/+.09mm (Female/Male) and M.L -2.0mm from Bregma, and -3.0mm ventral from dura corresponding to the Caudate Putamen region according to the mouse atlas of Paxinos and Franklin(72). Animals required for gene expression analysis were bilaterally injected with the same coordinates in the left hemisphere, but M.L. + 2.0 from bregma. Following the injection, animals were sutured with metal clips and subcutaneously injected with an antagonist mix; flumazenil (0,1mg/ml), naloxone (0,4 mg/ml), and atipamezole hydrochloride (5mg/ml) in 0,9% NaCl for wakening in a final dose of 10ml/kg (0,4-0,5ml/injection). The animals were injected with buprenorphine (Temgesic 0,3mg/ml, i.p.) after surgery for pain relief.

### Challenging beam test

Motor function and coordination were evaluated in the challenging beam test as previously described with small modifications(73). The mice have to transverse an increasingly narrowing beam from the widest part (frame 1) to the narrowest part (frame 4). Each frame is 25cm. The mice are trained for two days before testing. On test day, a grid is placed on top of the beam (leaving a 1cm space between the grid and the beam surface). Each animal undergoes five trial runs. A video camera is used to record all trials. All handling of the animals was done by the same experimenter. The videos are analyzed by an observer blind to the identity of the animals. The time to transverse the beam, the number of steps and errors are examined and the average of the five trials is presented. Frame 1 was excluded from the analysis due to high variability.

### Spontaneous activity in the cylinder test

The spontaneous activity levels and asymmetry of movements were measured in an adaptation of the cylinder test(73). The animals are placed in a glass cylinder and spontaneous activity is video recorded for a minimum of 3 min. The number of rearings (vertical activities) is simultaneously counted. A minimum of 20 rearings is set. In case the animals did not reach 20 rearings in 3 minutes, the experiment was prolonged until 20 rearings were reached with a maximum of 7 minutes. Each animal underwent one test. The number of left/right forelimb and hindlimb steps and time spent grooming (seconds) were analyzed within the 3 minutes. While the number of rearings and time to reach 20 rearings was analyzed for the whole video (<7minutes) if it exceeded the 3 minutes. It was not considered a step when the animal places a paw back of the cylinder bottom after a rearing. All analysis was done by an observer blind to the identity of the animals.

### Perfusion and post-fixation of the brain for immunohistochemistry

The animals were euthanized 1 month or 6 months after the injection of α-syn by an overdose of pentobarbital (400mg/ml, 1:10 dilution). On respiratory arrest, mice were perfused with cold physiological 0.9% NaCl solution through the ascending aorta, followed by paraformaldehyde (4%, ice-cold PFA, 0.1 NaPB, pH 7.4). The brains were removed and post-fixed in PFA solution for approx. 2 hours and then transferred to 25% sucrose solution (in 0.02M NaPB). The brains were sectioned on a freezing microtome (Brock and Michelsen, Thermo Fisher Scientific) into 40µm thick coronal sections and separated into serial sections; a series of 6 for the striatum, and a series of 4 for the SN. Sections were stored at -20°C in an anti-freeze solution.

### Immunohistochemistry

Immunohistochemistry was performed on free-floating sections. During the staining process, sections were washed multiple times in phosphate buffer (KPBS) between each incubation period. All incubation periods contained 0,25% Triton X-100 in KPBS. Shortly, the sections are quenched for 10 min in a solution of 3% hydrogen peroxide and 10% methanol, followed by one hour of blocking with the appropriate 5% serum. The sections were then incubated overnight at room temperature with the primary antibody (and 2,5% serum) (Sup. Table 1). The next day, sections were incubated for 2 hours with the appropriate biotinylated secondary antibody in 1% serum, followed by 1-hour incubation with avidin-biotin-peroxidase complex (vectorstain, ABCkit, Vector laboratories; PK-6100) in KPBS. Development was done with 3,3- diaminobenzidine (DAB) and reacted started with the appropriate concentration of H_2_O_2_. Afterward, sections were mounted on gelatin-coated glass slides and coverslipped. Iba-1 immunostained sections were counterstained with cresyl violet (0.5% solution) before coverslipping.

### Stereological analysis

Stereological estimation of tyrosine hydroxylase positive cells (TH^+^) in the SN was made by an observer blind to the identity of the animals. Sampling was done using the VIS software (Visiopharm A/S) and a bright field Leica DM6000B (Prior Scientific) microscope. Between 8-10 sections in a series of the SN were chosen to range from around -2.7mm and -4mm from bregma according to the mouse brain atlas(72). A super-image of the selected sections was taken using a 1.25X objective, and the borders of the SN were drawn in this magnification using the Region of interest (ROI) tool. The counting frame (56.89µm x 42.66µm) were placed randomly at the first counting by the VIS software and then systematically move through the whole ROI for each section. The step length was adjusted between 140-145µm for the contralateral side and between 120-135µm for the ipsilateral side, so approx. 100-150 TH+ cells were counted on both sides of the SN. The counting was conducted with a 40X objective. The total number of cells was calculated according to the optical fractionator formula and a coefficient of error (CE) < 0.10 was accepted.

Stereological estimation of Iba1^+^ cells in the SN and their morphology was made by an observer blind to the identity of the animals. The sampling software and approach to defining SN were the same as described for TH stereology and included both SN compacta and reticulata. The sampling frequency was adjusted between 140-150µm so a total of approx. 250 cells were counted in each side (ipsi and contra). The counting was performed using a 40x objective. The total number of cells was calculated by the optical fractionator formula and a coefficient of error (CE) < 0.10 was accepted.

### Densitometry

The optical density of the TH^+^ fibers in the striatum were measured at 6 different levels of rostro-caudal levels according to the mouse brain atlas(72) (+1.1; +0.74; +0.38; + 0.02; -0.34; -0.70 mm relative to bregma). The different brain sections were scanned using a densitometer (BioRad, GS-710), and the digital images were analyzed using the Fiji software using a greyscale. The optical density for each section was corrected with the nonspecific background measure from the corpus callosum. The quantification was made by an observer blind to the identity of the animals.

### Microscopy analysis

*MHCII+*: For quantification of ramified MHCII^+^ cells in the striatum, three representative sections in the striatum were a selection from each animal, located between: +1.18 and -0.94 mm from bregma according to the mouse brain atlas(72). The quantification was made using a Leica DMI600B microscope at 10X magnification. Due to the low expression and infiltration pattern of MHCII^+,^ a stereological counting approach was not chosen, and cells were manually quantified by an observer blind to the identity of the animals. MHCII^+^ cells at the site of injection and presence in the white matter were not quantified.

*MJF14+*: For quantification of MJF14 positive staining in the amygdala and SN an upright slide Scanner Microscope (Olympus VS120) was used at a magnification of 40x. In each animal, two sections for each area were selected. Enhanced Focal Imaging (EFI) photos were taken of these areas, with a depth (z-axis) of 20µm. The photos were analyzed with Fiji software using a protocol for unbiased quantification. First, the photos were converted to a binary 8-bit grayscale. The ROI was drawn in the photos and a histogram was obtained to receive information about the mode of the photo. The mode was multiplied by 0,87 (87%) to obtain the optimal threshold value when quantifying MJF14 staining. This optimal threshold value will distinguish specific staining from the background. Positive staining was quantified using the “analyze particles”-tool in Fiji. A lower cut-off of 5 pixels were set to avoid quantification of small unspecific staining. Circularity was set to 0-1 as aggregates are not uniform in morphology or circular. The same settings were applied to all animals. The percentage of the area (%Area fraction) occupied by the positive staining was calculated.

*p62+*: p62 staining was quantified in the amygdala and the SN. Two sections for each area were selected and 20x EFI photos with a depth (z-axis) of 20µm were taken (Olympus VS120). The same procedure as described for MJF14 was applied to quantified %Area fraction. The optimal threshold was determined to 80% of the mode, and the low-cut off pixels value to 20 pixels. Circularity was set to 0-1

*CD68+*: CD68 staining was quantified in striatal sections (three per animal), with images acquired at 20x magnification using EFI with a z-axis depth of 20µm (Olympus VS120). For the SN, two to three sections per animal were analyzed, and images were acquired at 20x with a z-axis depth of 22µm (11 layers with 2µm spacing) using NDP software (NanoZoomer 2.0HT, Hamamatsu). Quantification of the % area fraction was performed using the same procedure described for MJF14. The optimal threshold was determined to 80% of the mode, the low-cut off pixels value to 5 pixels, and circularity was defined between 0- 1. Values were average per section.

*pSer129+*: For pSer129+ one section with the frontal cortex and one section with the thalamus were selected, while two sections with the amygdala were selected. 20x EFI photos with a depth (z-axis) of 20µm were taken (Olympus VS120). For the SN, two sections were selected and 20x EDF images with a depth (z-axis) of 20µm were obtained (AxioScan 7. Zeiss). The same procedure as described for MJF14 was applied to quantify the %Area fraction. However, due to a more pronounced background staining in some samples, the optimal threshold was determined to be 80% of the mode in sections with less background, and 70% of the mode in animals with more background. The low-cut off pixels value was 5 pixels and circularity 0-1.

### Gene-panel assay (BioMark, Fluidigm)

For the gene-panel assay tissue was homogenized with the TissueLyser System (QIAGEN) for 3 min in an appropriate volume of PBS. RNA was isolated from the frontal cortex, striatum, and ventral midbrain from n=22 animals. Only male mice were used for the RNA analysis. (High RNA isolation Kit from Roche). Isolated RNA was dilution in elution buffer to a concentration of 12,5 ng/µl. In the following 2µl RNA was used from the diluted RNA. First, complementary DNA (cDNA) was produced from RNA. Samples were mixed with SuperScript III RT/Platinum Taq mix (Invitrogen, SuperScript® III platinum® one-Step qRT-PCR system) and 0.2x of all primers used in the study (Sup. Table 2) (TaqMan gene expression assayed, Applied Biosystems). The amplification with reverse transcription (RT-PCR) was performed using the cycling protocol: RT (incubated at 50°C for 15 min), Taq activation (95°C for 2 min), followed by 15 cycles of amplification (95°C at 15s and 60°C, at 4min). The cDNA was at diluted at least 1:10 and max 1:30 before real-time qPCR, which was performed in a 192.24 Dynamic Array Integrated Fluidic Circuits (Fluidigm) chip using the same primers as mentioned above. Technical duplicates were included for striatum samples. The following reagents were used for the qPCR reactions; TaqMan Gene expression master mix (Applied Biosystems), GE sample loading reagent (Fluidigm), and the pre-amplified cDNA. The primer mix was prepared with 2µl pf 20X TaqMan Gene expression assay (Applied Biosystems) and assay loading reagent (Fluidigm). The 192.24 Dynamic Array was primed in the IFC controller (Fluidigm) before loading. The sample mix, cDNA, and primer mix were loaded into the appropriate inlets and loading into the primed chip in the IFC controller. This setup allows each sample to combinate with each primer in separate reactions. The plate was placed in the BioMark PCR instrument (Fluidigm) and the standard protocol was followed. Data was required using the Fluidigm Real-Time PCR analysis software (Fluidigm). Data were normalized to the average of β-Act and GAPDH using the formula 2^ct(β-actin^ ^mRNA)-Ct(target^ ^mRNA)^. Failed Ct were removed from the analysis.

### Statistics

For statistical analysis of data GraphPad Prism 8 was utilized. For data with two variables (treatment and genotype) a Two-way ANOVA was applied followed by Sidak’s multiple comparison. For data with three variables (treatment, genotype, and brain hemisphere) a three-way ANOVA was conducted with matched values (Contralateral and ipsilateral). The three-way ANOVA was followed by an uncorrected Fishers LSD test and comparisons were done using Bonferroni corrections. Corrected p-values are displayed on the individual graphs. For further results regarding statistical analysis see Supplementary data.

## Acknowledgments

Funding support for the research covered in this article was provided by the Bjarne Saxhof Fund administered through the Danish Parkinson’s Foundation (M.R.-R.). The Lundbeck Foundation grants R223-2015-4222 and R248-2016-2518 for the Danish Research Institute of Translational Neuroscience-DANDRITE and R383-2022-180 (P.H.J. and H.G.). The work in the S.R.P. laboratory was funded by the Lundbeck Foundation (R359-2020-2287). I.K. was a recipient of a PhD fellowship from the Graduate School at the Health Faculty, Aarhus University, when performing the experiments described here.

## Credit authorship contribution statement

**IK** Writing – original draft, Investigation, Formal analysis. **LSR, SAF & JL:** Investigation, Formal analysis. **GUT:** technical support, **HG & PHJ**: provided resources, **SRP:** Conceptualization. **MRR:** Conceptualization, Project administration, Supervision, Investigation, Funding acquisition, Formal analysis, Data curation, Writing – review & editing, Writing – original draft. All authors reviewed and approved the final manuscript.

## Ethics declaration

M.R.-R. is Associate Editor of *npj Parkinson’s Disease*. M.R.-R. was not involved in the journal’s review of, or decisions related to, this manuscript. The remaining authors declare no competing financial or non-financial interests.

## Notes

### Competing Interest Statement

The authors have declared no competing interest.

### Summary of Updates

In this new revised version, we have performed additional experiments, new analysis, redesigned figures, and modified the text in line with the new data.

## References

1. W. Poewe et al., Parkinson disease. Nat Rev Dis Primers 3, 17013 (2017).

2. T. Tyson, J. A. Steiner, P. Brundin, Sorting out release, uptake and processing of alpha-synuclein during prion-like spread of pathology. Journal of neurochemistry 139 **Suppl 1**, 275–289 (2016).

3. T. Tyson, J. A. Steiner, P. Brundin, Sorting out release, uptake and processing of alpha-synuclein during prion-like spread of pathology. J Neurochem 139 **Suppl 1**, 275–289 (2016).

4. E. Emmanouilidou, K. Vekrellis, Exocytosis and Spreading of Normal and Aberrant α- Synuclein. Brain pathology (Zurich, Switzerland) 26, 398–403 (2016).

5. M. G. Tansey, M. Romero-Ramos, Immune system responses in Parkinson’s disease: Early and dynamic. Eur J Neurosci 49, 364–383 (2019).

6. T. Nagatsu, M. Mogi, H. Ichinose, A. Togari, Cytokines in Parkinson’s disease. J Neural Transm Suppl, 143–151 (2000).

7. A. Gerhard et al., In vivo imaging of microglial activation with [11C](R)-PK11195 PET in idiopathic Parkinson’s disease. Neurobiol Dis 21, 404–412 (2006).

8. K. Imamura et al., Distribution of major histocompatibility complex class II-positive microglia and cytokine profile of Parkinson’s disease brains. Acta neuropathologica 106, 518–526 (2003).

9. P. L. McGeer, S. Itagaki, B. E. Boyes, E. G. McGeer, Reactive microglia are positive for HLA-DR in the substantia nigra of Parkinson’s and Alzheimer’s disease brains. Neurology 38, 1285–1291 (1988).

10. S. A. Ferreira, M. Romero-Ramos, Microglia Response During Parkinson’s Disease: Alpha-Synuclein Intervention. Front Cell Neurosci 12, 247 (2018).

11. L. Fellner et al., Toll-like receptor 4 is required for α-synuclein dependent activation of microglia and astroglia. Glia 61, 349–360 (2013).

12. C. Kim et al., Neuron-released oligomeric α-synuclein is an endogenous agonist of TLR2 for paracrine activation of microglia. Nature communications 4, 1562 (2013).

13. N. Stefanova et al., Toll-Like Receptor 4 Promotes alpha-Synuclein Clearance and Survival of Nigral Dopaminergic Neurons. Am J Pathol 179, 954–963 (2011).

14. X. Su et al., Synuclein activates microglia in a model of Parkinson’s disease. Neurobiol Aging 29, 1690–1701 (2008).

15. H. J. Lee, J. E. Suk, E. J. Bae, S. J. Lee, Clearance and deposition of extracellular alpha-synuclein aggregates in microglia. Biochem Biophys Res Commun 372, 423–428 (2008).

16. J. Y. Park, S. R. Paik, I. Jou, S. M. Park, Microglial phagocytosis is enhanced by monomeric alpha-synuclein, not aggregated alpha-synuclein: implications for Parkinson’s disease. Glia 56, 1215–1223 (2008).

17. Q. Chen, L. Sun, Z. J. Chen, Regulation and function of the cGAS-STING pathway of cytosolic DNA sensing. Nature immunology 17, 1142–1149 (2016).

18. X. Li et al., Cyclic GMP-AMP synthase is activated by double-stranded DNA-induced oligomerization. Immunity 39, 1019–1031 (2013).

19. T. Abe, G. N. Barber, Cytosolic-DNA-mediated, STING-dependent proinflammatory gene induction necessitates canonical NF-kappaB activation through TBK1. Journal of virology 88, 5328–5341 (2014).

20. S. Liu et al., Phosphorylation of innate immune adaptor proteins MAVS, STING, and TRIF induces IRF3 activation. Science (New York, N.Y.) 347, aaa2630 (2015).

21. K. P. Hopfner, V. Hornung, Molecular mechanisms and cellular functions of cGAS-STING signalling. Nature reviews. Molecular cell biology 10.1038/s41580-020-0244-x (2020).

22. T. I. Kam et al., Poly(ADP-ribose) drives pathologic α-synuclein neurodegeneration in Parkinson’s disease. *Science (New York*, N.Y.) 362 (2018).

23. V. Vasquez et al., Chromatin-Bound Oxidized α-Synuclein Causes Strand Breaks in Neuronal Genomes in in vitro Models of Parkinson’s Disease. J Alzheimers Dis 60, S133–s150 (2017).

24. A. J. Schaser et al., Alpha-synuclein is a DNA binding protein that modulates DNA repair with implications for Lewy body disorders. Scientific reports 9, 10919 (2019).

25. T. Prabakaran et al., Attenuation of cGAS-STING signaling is mediated by a p62/SQSTM1- dependent autophagy pathway activated by TBK1. EMBO J 37 (2018).

26. X. Gui et al., Autophagy induction via STING trafficking is a primordial function of the cGAS pathway. Nature 567, 262–266 (2019).

27. A. Bayati, P. S. McPherson, Alpha-synuclein, autophagy-lysosomal pathway, and Lewy bodies: Mutations, propagation, aggregation, and the formation of inclusions. J Biol Chem 300, 107742 (2024).

28. K. L. Paumier et al., Intrastriatal injection of pre-formed mouse α-synuclein fibrils into rats triggers α-synuclein pathology and bilateral nigrostriatal degeneration. Neurobiology of disease 82, 185–199 (2015).

29. V. Sanchez-Guajardo, A. Annibali, P. H. Jensen, M. Romero-Ramos, alpha-Synuclein vaccination prevents the accumulation of parkinson disease-like pathologic inclusions in striatum in association with regulatory T cell recruitment in a rat model. J Neuropathol Exp Neurol 72, 624–645 (2013).

30. V. Sanchez-Guajardo, F. Febbraro, D. Kirik, M. Romero-Ramos, Microglia acquire distinct activation profiles depending on the degree of alpha-synuclein neuropathology in a rAAV based model of Parkinson’s disease. PLoS One 5, e8784 (2010).

31. D. J. Klionsky et al., Guidelines for the use and interpretation of assays for monitoring autophagy in higher eukaryotes. Autophagy 4, 151–175 (2008).

32. L. S. Reinert et al., Sensing of HSV-1 by the cGAS-STING pathway in microglia orchestrates antiviral defence in the CNS. Nat Commun 7, 13348 (2016).

33. Z. Tang et al., STING mediates lysosomal quality control and recovery through its proton channel function and TFEB activation in lysosomal storage disorders. Mol Cell 85, 1624–1639 e1625 (2025).

34. Y. Lu et al., Protective role of mitophagy on microglia-mediated neuroinflammatory injury through mtDNA-STING signaling in manganese-induced parkinsonism. J Neuroinflammation 22, 55 (2025).

35. K. C. Luk et al., Intracerebral inoculation of pathological alpha-synuclein initiates a rapidly progressive neurodegenerative alpha-synucleinopathy in mice. J Exp Med 209, 975–986 (2012).

36. J. T. Hinkle et al., STING mediates neurodegeneration and neuroinflammation in nigrostriatal α-synucleinopathy. Proceedings of the National Academy of Sciences of the United States of America 119, e2118819119 (2022).

37. K. C. Luk et al., Pathological alpha-synuclein transmission initiates Parkinson-like neurodegeneration in nontransgenic mice. Science 338, 949–953 (2012).

38. N. K. Polinski, A Summary of Phenotypes Observed in the In Vivo Rodent Alpha-Synuclein Preformed Fibril Model. J Parkinsons Dis 11, 1555–1567 (2021).

39. H. K. Chung, H. A. Ho, D. Perez-Acuna, S. J. Lee, Modeling alpha-Synuclein Propagation with Preformed Fibril Injections. J Mov Disord 13, 77–79 (2020).

40. K. C. Luk et al., Pathological α-synuclein transmission initiates Parkinson-like neurodegeneration in nontransgenic mice. *Science (New York*, N.Y.) 338, 949–953 (2012).

41. J. M. Froula et al., Defining α-synuclein species responsible for Parkinson’s disease phenotypes in mice. The Journal of biological chemistry 294, 10392–10406 (2019).

42. D. Liu et al., STING directly activates autophagy to tune the innate immune response. Cell Death & Differentiation 26, 1735–1749 (2019).

43. D. A. Chistiakov, M. C. Killingsworth, V. A. Myasoedova, A. N. Orekhov, Y. V. Bobryshev, CD68/macrosialin: not just a histochemical marker. Laboratory Investigation 97, 4–13 (2017).

44. A. Deczkowska et al., Disease-Associated Microglia: A Universal Immune Sensor of Neurodegeneration. Cell 173, 1073–1081 (2018).

45. S. A. Ferreira et al., Sex-dimorphic neuroprotective effect of CD163 in an alpha-synuclein mouse model of Parkinson’s disease. NPJ Parkinsons Dis 9, 164 (2023).

46. Y. Zhang, M. Zou, H. Wu, J. Zhu, T. Jin, The cGAS-STING pathway drives neuroinflammation and neurodegeneration via cellular and molecular mechanisms in neurodegenerative diseases. Neurobiol Dis 10.1016/j.nbd.2024.106710, 106710 (2024).

47. K. M. Crosse et al., Viperin binds STING and enhances the type-I interferon response following dsDNA detection. Immunol Cell Biol 99, 373–391 (2021).

48. S. Fruhwurth et al., TREM2 is down-regulated by HSV1 in microglia and involved in antiviral defense in the brain. Sci Adv 9, eadf5808 (2023).

49. M. Colonna, The biology of TREM receptors. Nat Rev Immunol 23, 580–594 (2023).

50. E. M. Szego et al., Constitutively active STING causes neuroinflammation and degeneration of dopaminergic neurons in mice. Elife 11 (2022).

51. E. L. Beatman et al., Alpha-Synuclein Expression Restricts RNA Viral Infections in the Brain. J Virol 90, 2767–2782 (2015).

52. N. Khanlou et al., Increased frequency of alpha-synuclein in the substantia nigra in human immunodeficiency virus infection. J Neurovirol 15, 131–138 (2009).

53. R. Marreiros et al., Disruption of cellular proteostasis by H1N1 influenza A virus causes α- synuclein aggregation. Proceedings of the National Academy of Sciences of the United States of America 117, 6741–6751 (2020).

54. B. Monogue et al., Alpha-synuclein supports type 1 interferon signalling in neurons and brain tissue. Brain : a journal of neurology 10.1093/brain/awac192 (2022).

55. F. Limanaqi et al., Alpha-synuclein dynamics bridge Type-I Interferon response and SARS-CoV-2 replication in peripheral cells. Biol Res 57, 2 (2024).

56. C. Kaufer et al., Microgliosis and neuronal proteinopathy in brain persist beyond viral clearance in SARS-CoV-2 hamster model. EBioMedicine 79, 103999 (2022).

57. P. Ejlerskov et al., Lack of Neuronal IFN-beta-IFNAR Causes Lewy Body- and Parkinson’s Disease-like Dementia. Cell 163, 324–339 (2015).

58. S. Zhao, A. D. Umpierre, L. J. Wu, Tuning neural circuits and behaviors by microglia in the adult brain. Trends Neurosci 47, 181–194 (2024).

59. B. Lee et al., Inflammatory and anti-inflammatory cytokines bidirectionally modulate amygdala circuits regulating anxiety. Cell 188, 2190–2202 e2115 (2025).

60. J. N. Kodangattil, G. Möddel, M. Müller, W. Weber, A. Gorji, The Inflammatory Chemokine CXCL10 Modulates Synaptic Plasticity and Neuronal Activity in the Hippocampus. European Journal of Inflammation 10, 311–328 (2012).

61. J. I. Jimenez-Loygorri et al., Mitophagy curtails cytosolic mtDNA-dependent activation of cGAS/STING inflammation during aging. Nat Commun 15, 830 (2024).

62. D. P. Narendra, R. J. Youle, The role of PINK1-Parkin in mitochondrial quality control. Nat Cell Biol 26, 1639–1651 (2024).

63. M. Borsche et al., Mitochondrial damage-associated inflammation highlights biomarkers in PRKN/PINK1 parkinsonism. Brain 143, 3041–3051 (2020).

64. P. Podlesniy et al., Mitochondrial DNA in CSF distinguishes LRRK2 from idiopathic Parkinson’s disease. Neurobiol Dis 94, 10–17 (2016).

65. R. Wang et al., Glucosylceramide accumulation in microglia triggers STING-dependent neuroinflammation and neurodegeneration in mice. Sci Signal 17, eadk8249 (2024).

66. M. Hegde, et al., Mitochondria-Targeted Oligomeric alpha-Synuclein Induces TOM40 Degradation and Mitochondrial Dysfunction in Parkinson’s Disease and Parkinsonism-Dementia of Guam. Res Sq 10.21203/rs.3.rs-3970470/v1 (2024).

67. C. Milanese et al., Activation of the DNA damage response in vivo in synucleinopathy models of Parkinson’s disease. Cell Death Dis 9, 818 (2018).

68. L. A. Volpicelli-Daley, K. C. Luk, V. M. Lee, Addition of exogenous alpha-synuclein preformed fibrils to primary neuronal cultures to seed recruitment of endogenous alpha-synuclein to Lewy body and Lewy neurite-like aggregates. Nat Protoc 9, 2135–2146 (2014).

69. N. K. Polinski et al., Best Practices for Generating and Using Alpha-Synuclein Pre-Formed Fibrils to Model Parkinson’s Disease in Rodents. Journal of Parkinson’s disease 8, 303–322 (2018).

70. M. B. Thomsen et al., PET imaging reveals early and progressive dopaminergic deficits after intra-striatal injection of preformed alpha-synuclein fibrils in rats. Neurobiol Dis 149, 105229 (2021).

71. H. Abdelmotilib et al., alpha-Synuclein fibril-induced inclusion spread in rats and mice correlates with dopaminergic Neurodegeneration. Neurobiol Dis 105, 84–98 (2017).

72. P. G. Franklin KBJ, Paxinos and Franklin’s the mouse brain in stereotaxic coordinates (Academic Press, Amsterdam, ed. fourth 2013).

73. S. M. Fleming, O. R. Ekhator, V. Ghisays, Assessment of sensorimotor function in mouse models of Parkinson’s disease. J Vis Exp 10.3791/50303 (2013).

