## Supplementary info for "Lack of functional STING modulates immunity but does not protect dopaminergic neurons in the alpha-synuclein preformed fibrils Parkinson’s Disease mouse model"

#### Supplementary Tables and figures

**Supplementary table 1: Antibodies used for immunohistochemistry**

| Antibody | Dilution | Type/Clone | Company |
|---|---|---|---|
| **TH** | 1:750 | Polyclonal | Millipore (AB152) |
| **MHCII** | 1:400 | Monoclonal/M5/114.15.2 | eBioscience (14-5321) |
| **MJF14** | 1:25000 | Monoclonal/MJFR14-6-4-2 | Abcam(ab209538) |
| **pSer129** | 1:3000 | Monoclonal/D1R1R | Cell signaling technology (23706) |
| **P62/SQSTM1** | 1:2000 | Polyclonal | Nordic Biosite (18420-1-AP) |
| **Iba-1** | 1:1000 | Polyclonal | Wako Fujifilm (019-19741) |
| **CD68** | 1:1000 | Monoclonal/FA-11 | Bio-Rad (MCA1957) |

**Supplementary table 2: Primers for biomarker assay (TaqMan gene expression assayed, Applied Biosystems).**

| mRNA target | Ref: |
|---|---|
| **CXCL10** | Mm00445235_m1 |
| **CXCL1** | Mm04207460_m1 |
| **CXCL2** | Mm00436450_m1 |
| **CCL2** | Mm00441242_m1 |
| **TNFα** | Mm00443260_g1 |
| **IFN-β** | Mm00439552_s1 |
| **MX1** | Mm00487796_m1 |
| **IFN-γ** | Mm01168134_m1 |
| **IL1-b** | Mm00434228_m1 |
| **IL6** | Mm00446190_m1 |
| **IL-10** | Mm01288386_m1 |
| **gCSF** | Mm00438334_m1 |
| **TREM2** | Mm04209424_g1 |
| **C1qa** | Mm00432142_m1 |
| **C4b** | Mm00437893_g1 |
| **TLR2** | Mm00442346_m1 |
| **TLR4** | Mm00445273_m1 |
| **TLR2** | Mm00442346_m1 |
| **Viperin/RSAD2** | Mm00491265_m1 |
| **PUMA** | Mm00519268_m1 |
| **β-actin** | Mm00607939_s1 |
| **GAPDH** | Mm99999915_g1 |

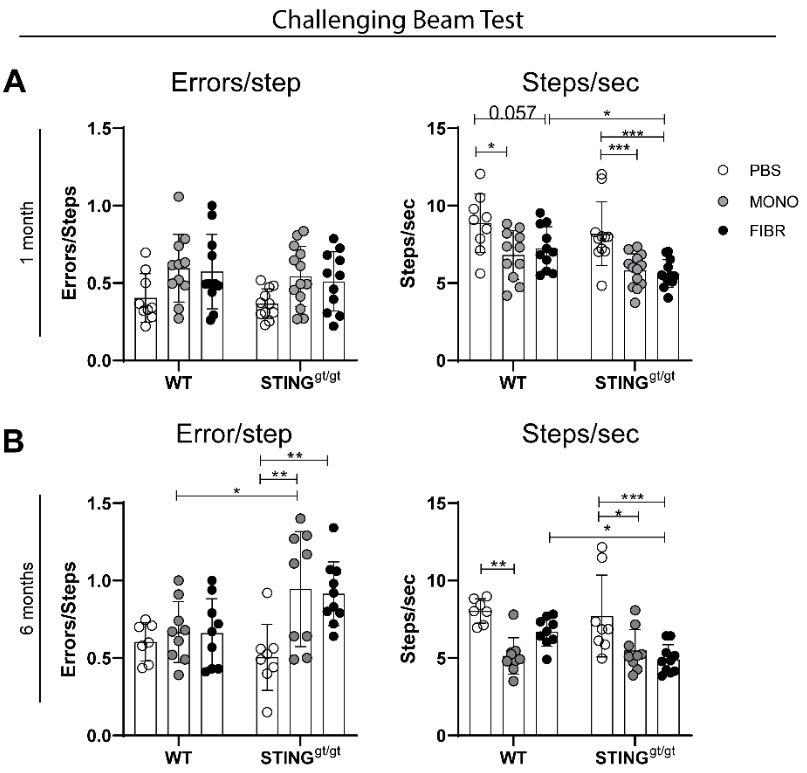

**Supplementary figure 1: Challenging beam test.** The challenging beam test was performed to evaluate motor behavior. At both 1-month **(A)** and 6-months post-injection **(B),** the number of errors/step and Steps/sec was calculated. n=7-10. For all graphs in the figure, a two-way ANOVA followed by Sidak's multiple comparisons was applied (test statistics in supplementary table 3). All values are mean +SD. *<p=0.05, **p≤0.01, ***p≤0.001, ****p≤0.0001.

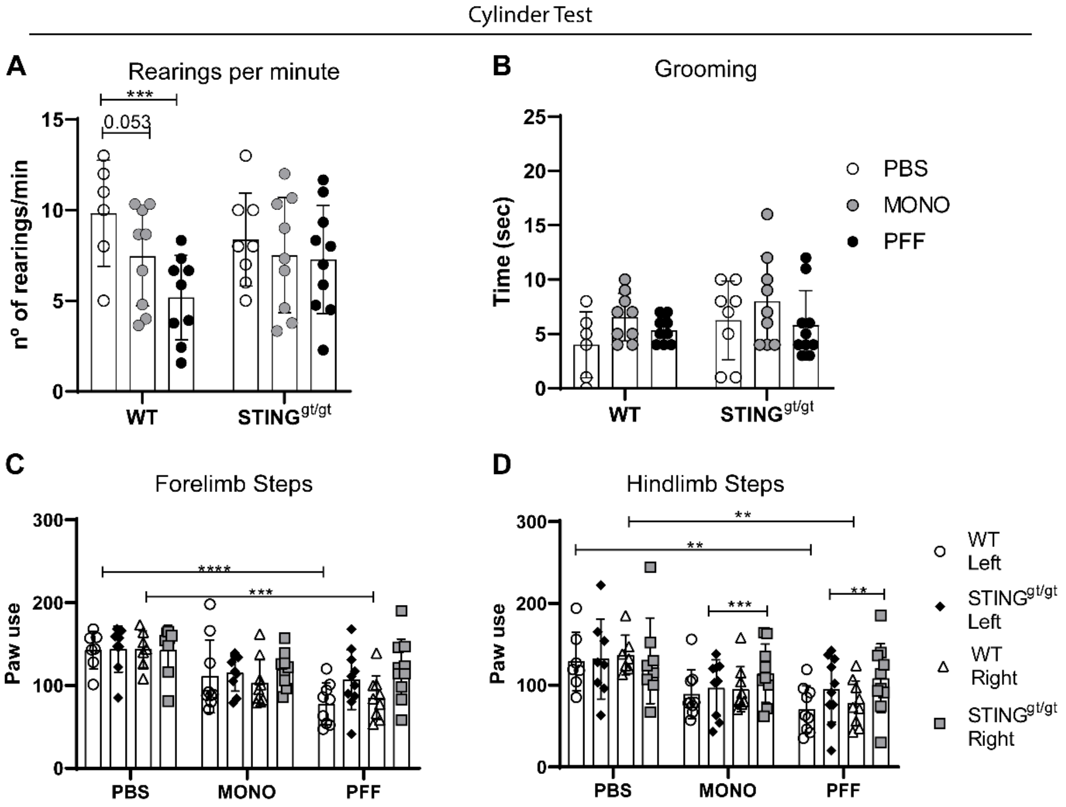

**Supplementary figure 2: the Spontaneous activity test.** At 6 months post-injection, the spontaneous activity in the cylinder test was performed to evaluate the motoric asymmetry. **A and B)** The number of rearings (A) and time spent grooming (B) were evaluated. **C and D)** The total paw use (steps) and the right and left forelimbs (C) and hindlimbs (D) were quantified. For A and B a two-way ANOVA followed by Sidak's multiple comparisons was applied (test statistics in supplementary table 3). For C and D). Three-way ANOVA with matched values followed by an uncorrected Fisher's LSD with Bonferroni correction applied. Corrected values are displayed on the graphs (*<p=0.05, **p≤0.01, ***p≤0.001, ****p≤0.0001). For all graphs, values are displayed as ± SD. n=6-10.

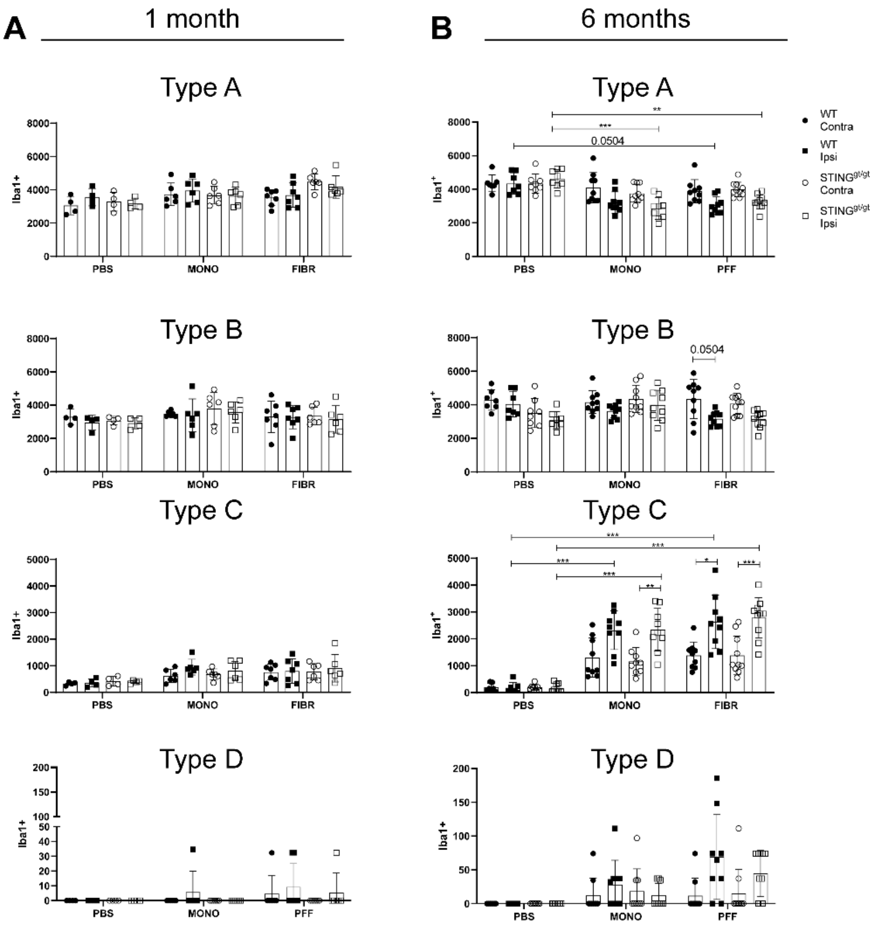

**Supplementary figure 3: Stereological quantification of iba1 morphology reveals a long-term activation of microglia. A. A and C)** Bar graphs illustrate the total number of either type A, B, C and D Iba1+ cells counted during the stereological quantification in the SN at 1 month p.i (B) and 6 months (C) post-injection . Three-way ANOVA with matched values followed by an uncorrected Fisher's LSD with Bonferroni correction applied. Corrected values are displayed on the graphs (*<p=0.05, **p≤0.01, ***p≤0.001, ****p≤0.0001). Data is displayed as ±SD, n=4-10. Note that total type D cells counted during the stereological quantification had a CE>0.01, thus no statistical test has been applied to the graphs displaying Type D Iba1$^{+}$ cells.

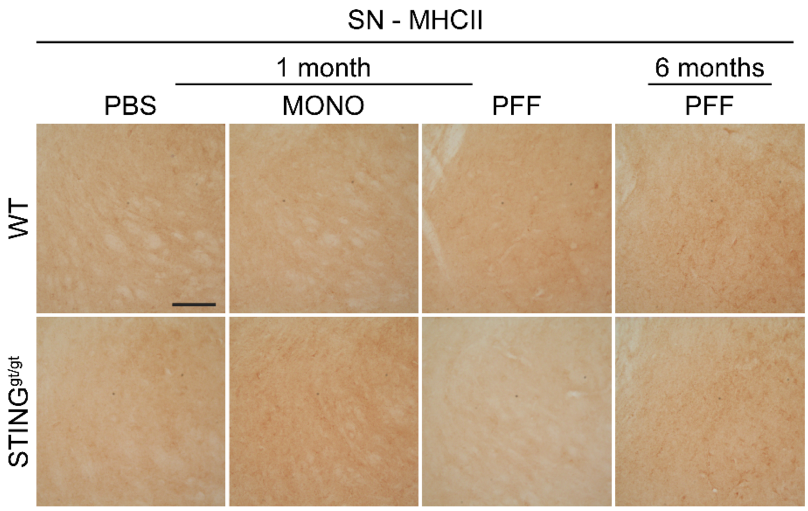

Supplementary Figure 4: Representative image of immune marker MHCII in the substantia nigra (SN) at 1 month and 6 months post-injection. No positive MHCII signaling was detected. Scalebar=100µm applies to all images.

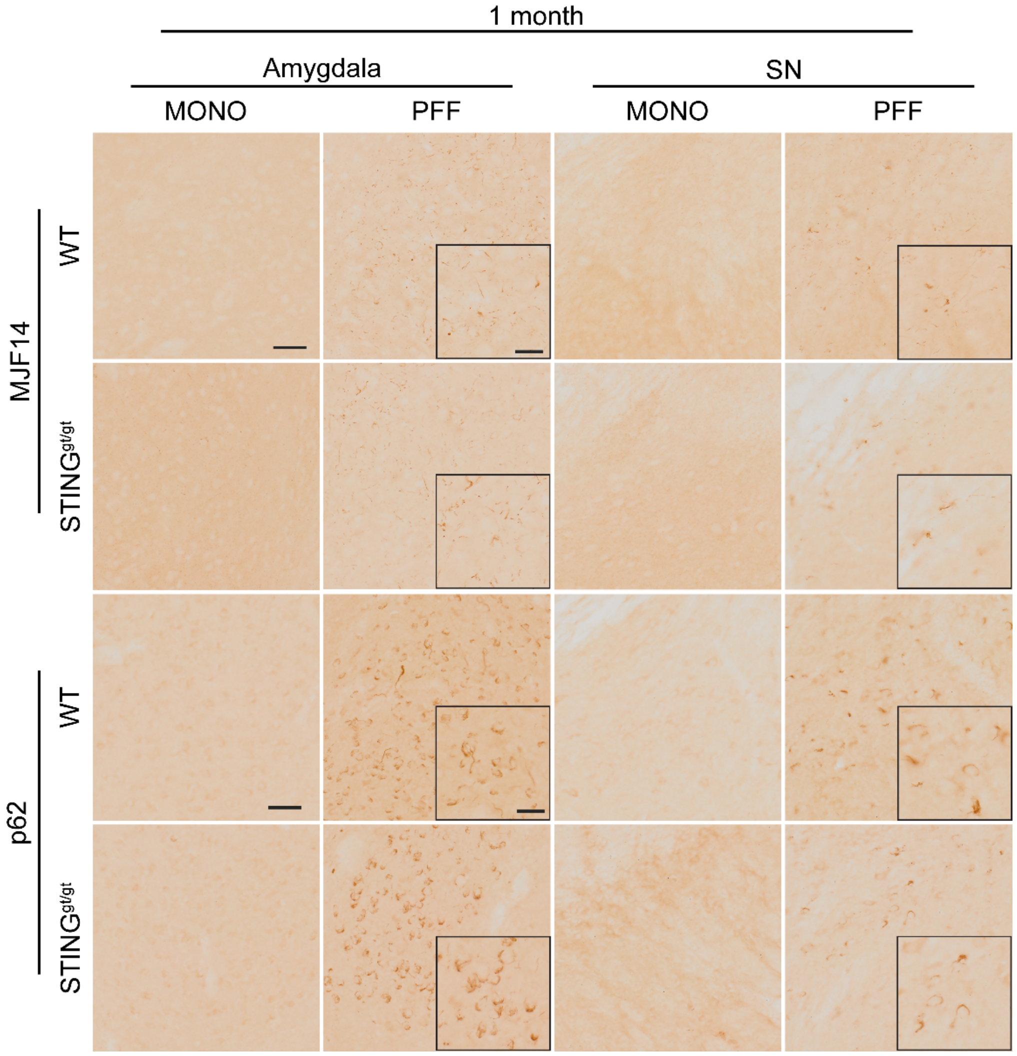

**Supplementary figure 5**: Photos of MJF14 and p62staining in amygdala and S. nigra at 1-month post-injection. As the area covered by p62 immunostaining in amygdala in PFF-mice was very variable (see main fig. 5), selected images show those animals with obvious p62 immuno-stained area (above average). Scalebar=50µm for all low magnification images and Scalebar=25µm for all inserts. One representative scalebar is shown for each area and marker.

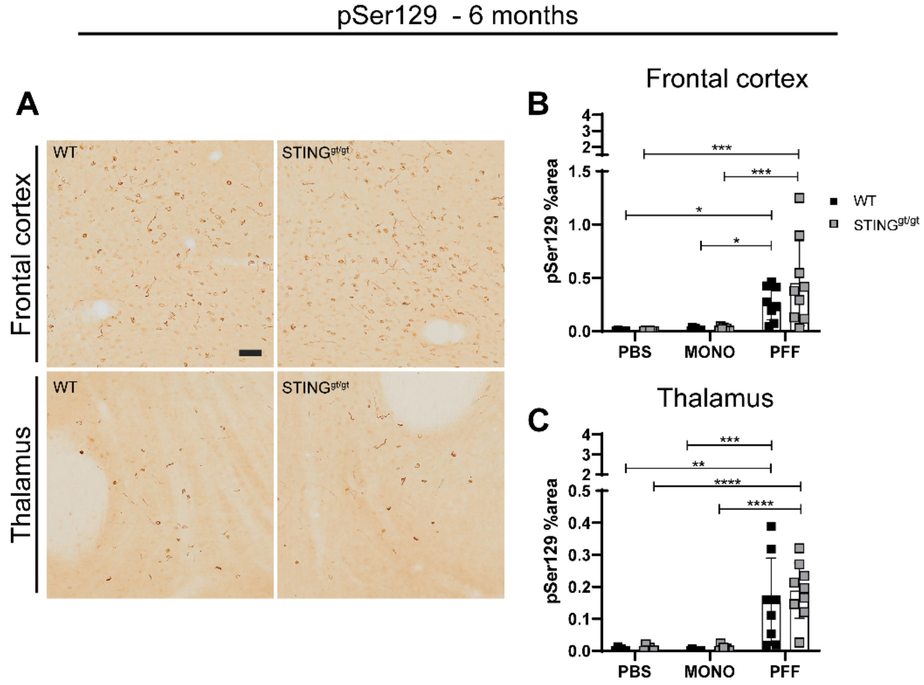

**Supplementary figure 6**: Injections of PFF lead to increase of pSer129-positive staining. **A)** Representative photos of pSer129 staining the frontal cortex and thalamus at 6 months post-injections. Scalebar=50μm applies to all images. **B, C )** Graphs show percentage of area covered by pser129 staining in PBS, MONO, and PFF-injected animals 6 months post-injected in the frontal cortex (B), and thalamus (C). Two-way ANOVA followed by Sidak's multiple comparisons (see further statistics in supplementary table 3). *<p=0.05, **p≤0.01, ***p≤0.001, ****p≤0.0001. All data are presented as mean ±SD. n= 6-10.

#### Supplementary table 3: 2-WAY ANOVA and 3-way ANOVA results

##### Figure 1- top panel

| **Challening beam Total time** | **ANOVA table** | **SS (Type III)** | **DF** | **MS** | **F (DFn, DFd)** | **P value** |
|---|---|---|---|---|---|---|
| | Interaction | 8.198 | 2 | 4.099 | F (2, 61) = 0,6668 | P=0,5170 |
| | Genotype | 25.83 | 1 | 25.83 | F (1, 61) = 4,202 | P=0,0447 |
| | Treatment | 145.3 | 2 | 72.63 | F (2, 61) = 11,82 | P<0,0001 |
| | Residual | 375 | 61 | 6.147 | | |
| **Challening beam Total steps** | **ANOVA table** | **SS (Type III)** | **DF** | **MS** | **F (DFn, DFd)** | **P value** |
| | Interaction | 3.121 | 2 | 1.561 | F (2, 61) = 1,544 | P=0,2218 |
| | Genotype | 5.703 | 1 | 5.703 | F (1, 61) = 5,642 | P=0,0207 |
| | Treatment | 9.978 | 2 | 4.989 | F (2, 61) = 4,935 | P=0,0103 |
| | Residual | 61.66 | 61 | 1.011 | | |
| **Challening beam Total Errors** | **ANOVA table** | **SS (Type III)** | **DF** | **MS** | **F (DFn, DFd)** | **P value** |
| | Interaction | 0.1243 | 2 | 0.06214 | F (2, 60) = 0,05073 | P=0,9506 |
| | Genotype | 0.3984 | 1 | 0.3984 | F (1, 60) = 0,3253 | P=0,5706 |
| | Treatment | 17.28 | 2 | 8.641 | F (2, 60) = 7,055 | P=0,0018 |
| | Residual | 73.49 | 60 | 1.225 | | |

##### Figure 1 - middle panel

| **Challening beam Total time** | **ANOVA table** | **SS (Type III)** | **DF** | **MS** | **F (DFn, DFd)** | **P value** |
|---|---|---|---|---|---|---|
| | Interaction | 34.34 | 2 | 17.17 | F (2, 45) = 2,064 | P=0,1388 |
| | Genotype | 17.42 | 1 | 17.42 | F (1, 45) = 2,095 | P=0,1547 |
| | Treatment | 211.4 | 2 | 105.7 | F (2, 45) = 12,71 | P<0,0001 |
| | Residual | 374.3 | 45 | 8.317 | | |
| **Challening beam Total steps** | **ANOVA table** | **SS (Type III)** | **DF** | **MS** | **F (DFn, DFd)** | **P value** |
| | Interaction | 5.647 | 2 | 2.823 | F (2, 46) = 1,499 | P=0,2340 |
| | Genotype | 0.158 | 1 | 0.158 | F (1, 46) = 0,08392 | P=0,7734 |
| | Treatment | 16.71 | 2 | 8.354 | F (2, 46) = 4,437 | P=0,0173 |
| | Residual | 86.61 | 46 | 1.883 | | |
| **Challening beam Total Errors** | **ANOVA table** | **SS (Type III)** | **DF** | **MS** | **F (DFn, DFd)** | **P value** |
| | Interaction | 14.85 | 2 | 7.427 | F (2, 46) = 3,851 | P=0,0284 |
| | Genotype | 7.767 | 1 | 7.767 | F (1, 46) = 4,027 | P=0,0507 |
| | Treatment | 16.15 | 2 | 8.077 | F (2, 46) = 4,188 | P=0,0213 |
| | Residual | 88.72 | 46 | 1.929 | | |

##### Figure 1 – lower panel

| **Forelimbs Assymetry: %left paw/Right paw** | **ANOVA table** | **SS (Type III)** | **DF** | **MS** | **F (DFn, DFd)** | **P value** |
|---|---|---|---|---|---|---|
| | Interaction | 43.42 | 2 | 21.71 | F (2, 46) = 0,4782 | P=0,6230 |
| | Genotype | 8.512 | 1 | 8.512 | F (1, 46) = 0,1875 | P=0,6670 |
| | Treatment | 970.8 | 2 | 485.4 | F (2, 46) = 10,69 | P=0,0002 |
| | Residual | 2088 | 46 | 45.4 | | |
| **Hindlimb Assymetry: %Left paw/Right paw** | **ANOVA table** | **SS (Type III)** | **DF** | **MS** | **F (DFn, DFd)** | **P value** |
| | Interaction | 817.5 | 2 | 408.7 | F (2, 46) = 2,298 | P=0,1119 |
| | Genotype | 14.16 | 1 | 14.16 | F (1, 46) = 0,07963 | P=0,7791 |
| | Treatment | 1035 | 2 | 517.6 | F (2, 46) = 2,910 | P=0,0646 |
| | Residual | 8182 | 46 | 177.9 | | |

##### Figure 2C

| **Total Iba1 cells** | **ANOVA table** | **SS** | **DF** | **MS** | **F (DFn, DFd)** | **P value** |
|---|---|---|---|---|---|---|
| | Treatment | 22346221 | 2 | 11173111 | F (2, 27) = 9,141 | P=0,0009 |
| | Genotype | 665548 | 1 | 665548 | F (1, 27) = 0,5445 | P=0,4669 |
| | Contra/Ipsi | 1097 | 1 | 1097 | F (1, 27) = 0,00327 | P=0,9548 |
| | Treatment x Genotype | 2394906 | 2 | 1197453 | F (2, 27) = 0,9797 | P=0,3884 |
| | Treatment x Contra/Ipsi | 264208 | 2 | 132104 | F (2, 27) = 0,3940 | P=0,6782 |
| | Genotype x Contra/Ipsi | 1119722 | 1 | 1119722 | F (1, 27) = 3,339 | P=0,0787 |
| | Treatment x Genotype x Contra/Ipsi | 11113 | 2 | 5557 | F (2, 27) = 0,01657 | P=0,9836 |
| | Subject | 33001382 | 27 | 1222273 | | |
| | Residual | 9053045 | 27 | 335298 | | |

#### Figure 2D

| | | | | | | | |
|---|---|---|---|---|---|---|---|
| **% of Iba1 (WT PBS: Contra x Ipsi)** | ANOVA table | SS | DF | MS | F (DFn, DFd) | P value | Repeate measure |
| | Hemisphere x Morphology | 161.6 | 3 | 53.86 | F (3, 12) = 5,557 | P=0,0126 | |
| | Hemisphere | 0 | 1 | 0 | F (1, 12) = 0,000 | P>0,9999 | |
| | Morphology | 16311 | 3 | 5437 | F (3, 12) = 3379 | P<0,0001 | |
| | Subject | 19.31 | 12 | 1.609 | F (12, 12) = 0,166( | P=0,9980 | |
| | Residual | 116.3 | 12 | 9.691 | | | |
| **% of Iba1 (SITNG PBS: Contra x Ipsi)** | ANOVA table | SS | DF | MS | F (DFn, DFd) | P value | Repeate measure |
| | Hemisphere x Morphology | 4.486 | 3 | 1.495 | F (3, 12) = 0,1269 | P=0,9423 | |
| | Hemisphere | 2.94 | 1 | 2.94 | F (1, 12) = 0,2496 | P=0,6264 | |
| | Morphology | 15385 | 3 | 5128 | F (3, 12) = 764,3 | P<0,0001 | |
| | Subject | 80.51 | 12 | 6.71 | F (12, 12) = 0,5695 | P=0,8287 | |
| | Residual | 141.4 | 12 | 11.78 | | | |
| **% of Iba1 (WT MONO: Contra x Ipsi)** | ANOVA table | SS | DF | MS | F (DFn, DFd) | P value | Repeate measure |
| | Hemisphere x Cell morphology | 95.46 | 3 | 31.82 | F (3, 20) = 1,251 | P=0,3179 | |
| | Hemisphere | 0.00003333 | 1 | 0.00003333 | F (1, 20) = 1,310e- | P=0,9991 | |
| | Cell morphology | 20134 | 3 | 6711 | F (3, 20) = 220,4 | P<0,0001 | |
| | Contra/Ipsi | 609.1 | 20 | 30.45 | F (20, 20) = 1,197 | P=0,3457 | |
| | Residual | 508.8 | 20 | 25.44 | | | |
| **% of Iba1 (WT PFF: Contra x Ipsi)** | **ANOVA table** | **SS** | **DF** | **MS** | **F (DFn, DFd)** | **P value** | Repeate measure |
| | Hemisphere x Cell morphology | 2.614 | 3 | 0.8713 | F (3, 24) = 0,0420( | P=0,9882 | |
| | Hemisphere | 7.143E-06 | 1 | 0.000007143 | F (1, 24) = 3,448e- | P=0,9995 | |
| | Cell morphology | 23235 | 3 | 7745 | F (3, 24) = 129,6 | P<0,0001 | |
| | Subject | 1435 | 24 | 59.78 | F (24, 24) = 2,886 | P=0,0060 | |
| | Residual | 497.1 | 24 | 20.71 | | | |
| **% of Iba1 (STING MONO: Contra x Ipsi)** | **ANOVA table** | **SS** | **DF** | **MS** | **F (DFn, DFd)** | **P value** | Repeate measure |
| | Hemisphere x Cell morphology | 16.97 | 3 | 5.656 | F (3, 20) = 0,2804 | P=0,8389 | |
| | Hemisphere | 8.333E-06 | 1 | 0.000008333 | F (1, 20) = 4,131e- | P=0,9995 | |
| | Cell morphology | 20471 | 3 | 6824 | F (3, 20) = 151,5 | P<0,0001 | |
| | Contra/Ipsi | 900.6 | 20 | 45.03 | F (20, 20) = 2,232 | P=0,0400 | |
| | Residual | 403.5 | 20 | 20.17 | | | |
| **% of Iba1 (STING FFF: Contra x Ipsi)** | **ANOVA table** | **SS** | **DF** | **MS** | **F (DFn, DFd)** | **P value** | Repeate measure |
| | Hemisphere x Cell morphology | 16.58 | 3 | 5.527 | F (3, 20) = 0,3408 | P=0,7961 | |
| | Hemisphere | 2.083E-06 | 1 | 0.000002083 | F (1, 20) = 1,285e- | P=0,9997 | |
| | Cell morphology | 21000 | 3 | 7000 | F (3, 20) = 177,2 | P<0,0001 | |
| | Contra/Ipsi | 790 | 20 | 39.5 | F (20, 20) = 2,436 | P=0,0265 | |
| | Residual | 324.3 | 20 | 16.22 | | | |
| **% of Iba1 (Ipsi PFF: WT x STING)** | **ANOVA table** | **SS** | **DF** | **MS** | **F (DFn, DFd)** | **P value** | |
| | Interaction | 149.9 | 3 | 49.96 | F (3, 44) = 1,546 | P=0,2159 | |
| | Genotype | 3.432 | 1 | 3.432 | F (1, 44) = 0,1062 | P=0,7460 | |
| | Morphology | 20795 | 3 | 6932 | F (3, 44) = 214,6 | P<0,0001 | |
| | Residual | 1421 | 44 | 32.31 | | | |
| **% of Iba1 (Ipsi MONO: WT x STING)** | **ANOVA table** | **SS** | **DF** | **MS** | **F (DFn, DFd)** | **P value** | |
| | Interaction | 74.49 | 3 | 24.83 | F (3, 40) = 0,6916 | P=0,5626 | |
| | Genotype | 8.333E-06 | 1 | 0.000008333 | F (1, 40) = 2,321e- | P=0,9996 | |
| | Morphology | 19192 | 3 | 6397 | F (3, 40) = 178,2 | P<0,0001 | |
| | Residual | 1436 | 40 | 35.91 | | | |
| **% of Iba1 (Contra PFF: WT x STING)** | **ANOVA table** | **SS** | **DF** | **MS** | **F (DFn, DFd)** | **P value** | |
| | Interaction | 132.2 | 3 | 44.08 | F (3, 44) = 1,290 | P=0,2896 | |
| | Genotype | 7.738E-06 | 1 | 0.000007738 | F (1, 44) = 2,265e- | P=0,9996 | |
| | Morphology | 22421 | 3 | 7474 | F (3, 44) = 218,8 | P<0,0001 | |
| | Residual | 1503 | 44 | 34.16 | | | |
| **% of Iba1 (Contra MONO: WT x STING)** | **ANOVA table** | **SS** | **DF** | **MS** | **F (DFn, DFd)** | **P value** | |
| | Interaction | 14.34 | 3 | 4.78 | F (3, 40) = 0,1940 | P=0,8999 | |
| | Genotype | 0 | 1 | 0 | F (1, 40) = 0,000 | P>0,9999 | |
| | Morphology | 21436 | 3 | 7145 | F (3, 40) = 289,9 | P<0,0001 | |
| | Residual | 985.8 | 40 | 24.64 | | | |
| **%Iba1 (Contra PBS: WT x STING)** | **ANOVA table** | **SS** | **DF** | **MS** | **F (DFn, DFd)** | **P value** | |
| | Interaction | 54.74 | 3 | 18.25 | F (3, 24) = 1,790 | P=0,1759 | |
| | Genotype | 0.1985 | 1 | 0.1985 | F (1, 24) = 0,01947 | P=0,8902 | |
| | Morphology | 16059 | 3 | 5353 | F (3, 24) = 525,3 | P<0,0001 | |
| | Residual | 244.6 | 24 | 10.19 | | | |
| **%Iba1 (Ipsi PBS: WT x STING)** | **ANOVA table** | **SS** | **DF** | **MS** | **F (DFn, DFd)** | **P value** | |
| | Interaction | 52.28 | 3 | 17.43 | F (3, 24) = 3,704 | P=0,0254 | |
| | Genotype | 1.611 | 1 | 1.611 | F (1, 24) = 0,3424 | P=0,5639 | |
| | Morphology | 15697 | 3 | 5232 | F (3, 24) = 1112 | P<0,0001 | |
| | Residual | 112.9 | 24 | 4.705 | | | |

#### Figure 2E

| **Total Iba1 cells** | **ANOVA table** | **SS** | **DF** | **MS** | **F (DFn, DFd)** | **P value** |
|---|---|---|---|---|---|---|
| | Treatment | 98452 | 1 | 98452 | F (1, 33) = 0,1202 | P=0,7310 |
| | Genotype | 9047 | 1 | 9047 | F (1, 33) = 0,01105 | P=0,9169 |
| | contra/ipsi | 1557671 | 1 | ##### | F (1, 33) = 3,013 | P=0,0919 |
| | Treatment x Genotype | 535229 | 1 | ##### | F (1, 33) = 0,6536 | P=0,4246 |
| | Treatment x contra/ipsi | 47351 | 1 | 47351 | F (1, 33) = 0,09159 | P=0,7641 |
| | Genotype x contra/ipsi | 294399 | 1 | ##### | F (1, 33) = 0,5694 | P=0,4558 |
| | Treatment x Genotype x contra/ipsi | 1973 | 1 | 1973 | F (1, 33) = 0,00381( | P=0,9511 |
| | Subject | 27023093 | 33 | ##### | | |
| | Residual | 17061144 | 33 | ##### | | |

#### Figure 2F

| | | | | | | | |
|---|---|---|---|---|---|---|---|
| **% of Iba1 (WT PBS: Contra x Ipsi)** | ANOVA table | SS | DF | MS | F (DFn, DFd) | P value | Repeate measure |
| | Hemisphere x Morphology | 13.64 | 3 | 4.548 | F (3, 24) = 0,3083 | P=0,8191 | |
| | Hemisphere | 0.00002857 | 1 | 3E-05 | F (1, 24) = 1,937e-0 | P=0,9989 | |
| | Morphology | 32245 | 3 | 10748 | F (3, 24) = 493,9 | P<0,0001 | |
| | Subject | 522.3 | 24 | 21.76 | F (24, 24) = 1,475 | P=0,1737 | |
| | Residual | 354 | 24 | 14.75 | | | |
| **% of Iba1 (STING PBS: Contra x Ipsi)** | ANOVA table | SS | DF | MS | F (DFn, DFd) | P value | Repeate measure |
| | Hemisphere x Morphology | 167.5 | 3 | 55.85 | F (3, 28) = 5,824 | P=0,0032 | |
| | Hemisphere | 2.827 | 1 | 2.827 | F (1, 28) = 0,2947 | P=0,5915 | |
| | Morphology | 39042 | 3 | 13014 | F (3, 28) = 284,7 | P<0,0001 | |
| | Subject | 1280 | 28 | 45.71 | F (28, 28) = 4,767 | P<0,0001 | |
| | Residual | 268.5 | 28 | 9.59 | | | |
| **% of Iba1 (WT MONO: Contra x Ipsi)** | ANOVA table | SS | DF | MS | F (DFn, DFd) | P value | Repeate measure |
| | Hemisphere x Cell morphology | 695.2 | 3 | 231.7 | F (3, 32) = 14,82 | P<0,0001 | |
| | Hemisphere | 0.06722 | 1 | 0.067 | F (1, 32) = 0,004299 | P=0,9481 | |
| | Cell morphology | 20054 | 3 | 6685 | F (3, 32) = 120,0 | P<0,0001 | |
| | contra/ipsi | 1782 | 32 | 55.69 | F (32, 32) = 3,561 | P=0,0003 | |
| | Residual | 500.4 | 32 | 15.64 | | | |
| **% of Iba1 (WT PFF: Contra x Ipsi)** | **ANOVA table** | **SS** | **DF** | **MS** | **F (DFn, DFd)** | **P value** | Repeate measure |
| | Hemisphere x Cell morphology | 1575 | 3 | 525 | F (3, 32) = 19,37 | P<0,0001 | |
| | Hemisphere | 8.208 | 1 | 8.208 | F (1, 32) = 0,3028 | P=0,5859 | |
| | Cell morphology | 18490 | 3 | 6163 | F (3, 32) = 111,5 | P<0,0001 | |
| | Contra/Ipsi | 1769 | 32 | 55.27 | F (32, 32) = 2,039 | P=0,0239 | |
| | Residual | 867.3 | 32 | 27.1 | | | |
| **% of Iba1 (STING MONO: Contra x Ipsi)** | **ANOVA table** | **SS** | **DF** | **MS** | **F (DFn, DFd)** | **P value** | Repeate measure |
| | Hemisphere x Cell morphology | 1267 | 3 | 422.2 | F (3, 32) = 26,55 | P<0,0001 | |
| | Hemisphere | 5.556E-06 | 1 | 6E-06 | F (1, 32) = 3,493e-0 | P=0,9995 | |
| | Cell morphology | 20860 | 3 | 6953 | F (3, 32) = 146,9 | P<0,0001 | |
| | Contra/Ipsi | 1515 | 32 | 47.34 | F (32, 32) = 2,977 | P=0,0014 | |
| | Residual | 508.9 | 32 | 15.9 | | | |
| **% of Iba1 (STING FFF: Contra x Ipsi)** | **ANOVA table** | **SS** | **DF** | **MS** | **F (DFn, DFd)** | **P value** | Repeate measure |
| | Hemisphere x Cell morphology | 1815 | 3 | 605 | F (3, 36) = 34,04 | P<0,0001 | |
| | Hemisphere | 2.019E-27 | 1 | 2E-27 | F (1, 36) = 1,136e-0 | P>0,9999 | |
| | Cell morphology | 19829 | 3 | 6610 | F (3, 36) = 193,2 | P<0,0001 | |
| | Contra/Ipsi | 1232 | 36 | 34.21 | F (36, 36) = 1,925 | P=0,0265 | |
| | Residual | 639.8 | 36 | 17.77 | | | |
| **% of Iba1 (Ipsi PFF: WT x STING)** | **ANOVA table** | **SS** | **DF** | **MS** | **F (DFn, DFd)** | **P value** | |
| | ANOVA table | SS (Type III) | DF | MS | F (DFn, DFd) | P value | |
| | Interaction | 20.67 | 3 | 6.889 | F (3, 68) = 0,2137 | P=0,8866 | |
| | Genotype | 1.345E-28 | 1 | 1E-28 | F (1, 68) = 4,173e-0 | P>0,9999 | |
| | Cell morphology | 15369 | 3 | 5123 | F (3, 68) = 158,9 | P<0,0001 | |
| **% of Iba1 (Ipsi MONO: WT x STING)** | **ANOVA table** | **SS** | **DF** | **MS** | **F (DFn, DFd)** | **P value** | |
| | Interaction | 98.48 | 3 | 32.83 | F (3, 64) = 0,9276 | P=0,4326 | |
| | Genotype | 1.291 | 1 | 1.291 | F (1, 64) = 0,03647 | P=0,8492 | |
| | Cell morphology | 17303 | 3 | 5768 | F (3, 64) = 163,0 | P<0,0001 | |
| | Residual | 2265 | 64 | 35.39 | | | |
| **% of Iba1 (Contra PFF: WT x STING)** | **ANOVA table** | **SS** | **DF** | **MS** | **F (DFn, DFd)** | **P value** | |
| | Interaction | 33.25 | 3 | 11.08 | F (3, 68) = 0,3255 | P=0,8069 | |
| | Genotype | 8.64 | 1 | 8.64 | F (1, 68) = 0,2538 | P=0,6160 | |
| | Cell morphology | 26203 | 3 | 8734 | F (3, 68) = 256,6 | P<0,0001 | |
| | Residual | 2315 | 68 | 34.04 | | | |
| **% of Iba1 (Contra MONO: WT x STING)** | **ANOVA table** | **SS** | **DF** | **MS** | **F (DFn, DFd)** | **P value** | |
| | Interaction | 122.6 | 3 | 40.86 | F (3, 64) = 1,281 | P=0,2885 | |
| | Genotype | 1.954 | 1 | 1.954 | F (1, 64) = 0,06124 | P=0,8053 | |
| | Cell morphology | 25351 | 3 | 8450 | F (3, 64) = 264,9 | P<0,0001 | |
| | Residual | 2042 | 64 | 31.9 | | | |
| **%Iba1 (Contra PBS: WT x STING)** | **ANOVA table** | **SS** | **DF** | **MS** | **F (DFn, DFd)** | **P value** | |
| | Interaction | 190.3 | 3 | 63.43 | F (3, 52) = 2,997 | P=0,0390 | |
| | Genotype | 1.071E-06 | 1 | 1E-06 | F (1, 52) = 5,062e-0 | P=0,9998 | |
| | Morphologt | 34331 | 3 | 11444 | F (3, 52) = 540,7 | P<0,0001 | |
| | Residual | 1101 | 52 | 21.17 | | | |
| **%Iba1 (Ipsi PBS: WT x STING)** | **ANOVA table** | **SS** | **DF** | **MS** | **F (DFn, DFd)** | **P value** | |
| | Interaction | 500 | 3 | 166.7 | F (3, 52) = 6,545 | P=0,0008 | |
| | Genotype | 2.66 | 1 | 2.66 | F (1, 52) = 0,1044 | P=0,7479 | |
| | Morphology | 35984 | 3 | 11995 | F (3, 52) = 471,0 | P<0,0001 | |
| | Residual | 1324 | 52 | 25.46 | | | |

#### Figure 3B

| **MHCII Striatum Average ramified MHCII** | **ANOVA table** | **SS (Type III)** | **DF** | **MS** | **F (DFn, DFd)** | **P value** |
|---|---|---|---|---|---|---|
| | Interaction | 55,4 | 1 | 55,4 | F (1, 20) = 0,04929 | P=0,8266 |
| | Genotype | 32,96 | 1 | 32,96 | F (1, 20) = 0,02932 | P=0,8658 |
| | Treatment | 14562 | 1 | 14562 | F (1, 20) = 12,96 | P=0,0018 |
| | Residual | 22481 | 20 | 1124 | | |

#### Figure 4B

| CD68 striatum %area | ANOVA table | SS (Type III) | DF | MS | F (DFn, DFd) | P value |
|---|---|---|---|---|---|---|
| | Interaction | 4,474 | 2 | 2,237 | F (2, 28) = 9,942 | P=0,0005 |
| | Treatment | 6,999 | 2 | 3,499 | F (2, 28) = 15,55 | P<0,0001 |
| | Genotype | 1,411 | 1 | 1,411 | F (1, 28) = 6,270 | P=0,0184 |
| | Residual | 6,3 | 28 | 0,225 | | |

#### Figure 4C

| CD68 striatum %area | ANOVA table | SS (Type III) | DF | MS | F (DFn, DFd) | P value |
|---|---|---|---|---|---|---|
| | Interaction | 0,09374 | 2 | 0,047 | F (2, 46) = 2,234 | P=0,1186 |
| | Treatment | 3,431 | 2 | 1,715 | F (2, 46) = 81,77 | P<0,0001 |
| | Genotype | 0,0001341 | 1 | 1E-04 | F (1, 46) = 0,006392 | P=0,9366 |
| | Residual | 0,965 | 46 | 0,021 | | |

#### Figure 4E

| CD68 SN %area | ANOVA table | SS (Type III) | DF | MS | F (DFn, DFd) | P value |
|---|---|---|---|---|---|---|
| | Interaction | 0.3017 | 2 | 0.1509 | F (2, 28) = 0,9934 | P=0,3830 |
| | Treatment | 0.2675 | 2 | 0.1338 | F (2, 28) = 0,8808 | P=0,4256 |
| | Genotype | 0.2061 | 1 | 0.2061 | F (1, 28) = 1,357 | P=0,2539 |
| | Residual | 4.252 | 28 | 0.1519 | | |

#### Figure 4F

| CD68 SN %area | ANOVA table | SS (Type III) | DF | MS | F (DFn, DFd) | P value |
|---|---|---|---|---|---|---|
| | Interaction | 0.2582 | 2 | 0.1291 | F (2, 35) = 1,486 | P=0,2402 |
| | Treatment | 1.768 | 2 | 0.8839 | F (2, 35) = 10,17 | P=0,0003 |
| | Genotype | 0.0124 | 1 | 0.0124 | F (1, 35) = 0,1427 | P=0,7079 |
| | Residual | 3.041 | 35 | 0.08689 | | |

### Figure 5A Prefrontal cortex

| | | | | | | |
|---|---|---|---|---|---|---|
| **TNF-a** | **ANOVA table** | **SS (Type III)** | **DF** | **MS** | **F (DFn, DFd)** | **P value** |
| | Interaction | 1.093E-07 | 1 | 1.093E-07 | F (1, 11) = 2,105 | P=0,1747 |
| | Genotype | 9.405E-08 | 1 | 9.405E-08 | F (1, 11) = 1,812 | P=0,2054 |
| | Treatment | 7.855E-08 | 1 | 7.855E-08 | F (1, 11) = 1,513 | P=0,2443 |
| | Residual | 5.71E-07 | 11 | 5.191E-08 | | |
| **IFN-b** | **ANOVA table** | **SS (Type III)** | **DF** | **MS** | **F (DFn, DFd)** | **P value** |
| | Interaction | 4.15E-09 | 1 | 4.15E-09 | F (1, 12) = 2,860 | P=0,1166 |
| | Genotype | 1.382E-11 | 1 | 1.382E-11 | F (1, 12) = 0,00952 | P=0,9239 |
| | Treatment | 6.781E-10 | 1 | 6.781E-10 | F (1, 12) = 0,4673 | P=0,5072 |
| | Residual | 1.741E-08 | 12 | 1.451E-09 | | |
| **IL1-b** | **ANOVA table** | **SS (Type III)** | **DF** | **MS** | **F (DFn, DFd)** | **P value** |
| | Interaction | 7.744E-11 | 1 | 7.744E-11 | F (1, 13) = 0,1450 | P=0,7095 |
| | Genotype | 3.6E-11 | 1 | 3.6E-11 | F (1, 13) = 0,06740 | P=0,7992 |
| | Treatment | 1.318E-09 | 1 | 1.318E-09 | F (1, 13) = 2,467 | P=0,1403 |
| | Residual | 6.944E-09 | 13 | 5.341E-10 | | |
| **IL-6** | **ANOVA table** | **SS (Type III)** | **DF** | **MS** | **F (DFn, DFd)** | **P value** |
| | Interaction | 4.277E-10 | 1 | 4.277E-10 | F (1, 16) = 1,107 | P=0,3084 |
| | Genotype | 1.632E-09 | 1 | 1.632E-09 | F (1, 16) = 4,225 | P=0,0565 |
| | Treatment | 2.9E-10 | 1 | 2.9E-10 | F (1, 16) = 0,7505 | P=0,3991 |
| | Residual | 6.182E-09 | 16 | 3.864E-10 | | |
| **CXCL10** | **ANOVA table** | **SS (Type III)** | **DF** | **MS** | **F (DFn, DFd)** | **P value** |
| | Interaction | 8.085E-08 | 1 | 8.085E-08 | F (1, 15) = 1,174 | P=0,2958 |
| | Genotype | 2.31E-09 | 1 | 2.31E-09 | F (1, 15) = 0,03353 | P=0,8572 |
| | Treatment | 2.254E-07 | 1 | 2.254E-07 | F (1, 15) = 3,272 | P=0,0906 |
| | Residual | 1.033E-06 | 15 | 6.89E-08 | | |
| **Mx1** | **ANOVA table** | **SS (Type III)** | **DF** | **MS** | **F (DFn, DFd)** | **P value** |
| | Interaction | 2.287E-09 | 1 | 2.287E-09 | F (1, 15) = 2,564 | P=0,1302 |
| | Genotype | 1.849E-09 | 1 | 1.849E-09 | F (1, 15) = 2,073 | P=0,1704 |
| | Treatment | 2.808E-11 | 1 | 2.808E-11 | F (1, 15) = 0,03149 | P=0,8615 |
| | Residual | 1.338E-08 | 15 | 8.919E-10 | | |
| **Viperin** | **ANOVA table** | **SS (Type III)** | **DF** | **MS** | **F (DFn, DFd)** | **P value** |
| | Interaction | 7.947E-10 | 1 | 7.947E-10 | F (1, 17) = 0,05542 | P=0,8167 |
| | Genotype | 1.756E-10 | 1 | 1.756E-10 | F (1, 17) = 0,01225 | P=0,9132 |
| | Treatment | 1.905E-08 | 1 | 1.905E-08 | F (1, 17) = 1,328 | P=0,2651 |
| | Residual | 2.438E-07 | 17 | 1.434E-08 | | |
| **CCL2** | **ANOVA table** | **SS (Type III)** | **DF** | **MS** | **F (DFn, DFd)** | **P value** |
| | Interaction | 1.064E-09 | 1 | 1.064E-09 | F (1, 14) = 3,758 | P=0,0730 |
| | Genotype | 1.321E-09 | 1 | 1.321E-09 | F (1, 14) = 4,667 | P=0,0486 |
| | Treatment | 6.847E-12 | 1 | 6.847E-12 | F (1, 14) = 0,02419 | P=0,8786 |
| | Residual | 3.963E-09 | 14 | 2.831E-10 | | |
| **TLR4** | **ANOVA table** | **SS (Type III)** | **DF** | **MS** | **F (DFn, DFd)** | **P value** |
| | Interaction | 3.016E-09 | 1 | 3.016E-09 | F (1, 15) = 12,43 | P=0,0031 |
| | Genotype | 2.311E-11 | 1 | 2.311E-11 | F (1, 15) = 0,09527 | P=0,7618 |
| | Treamtent | 1.29E-10 | 1 | 1.29E-10 | F (1, 15) = 0,5320 | P=0,4770 |
| | Residual | 3.638E-09 | 15 | 2.425E-10 | | |
| **TLR2** | **ANOVA table** | **SS (Type III)** | **DF** | **MS** | **F (DFn, DFd)** | **P value** |
| | Interaction | 8.684E-09 | 1 | 8.684E-09 | F (1, 15) = 9,633 | P=0,0073 |
| | Row Factor | 3.276E-09 | 1 | 3.276E-09 | F (1, 15) = 3,634 | P=0,0760 |
| | Column Factor | 1.636E-08 | 1 | 1.636E-08 | F (1, 15) = 18,15 | P=0,0007 |
| | Residual | 1.352E-08 | 15 | 9.014E-10 | | |
| **C1q** | **ANOVA table** | **SS (Type III)** | **DF** | **MS** | **F (DFn, DFd)** | **P value** |
| | Interaction | 0.00000266 | 1 | 0.00000266 | F (1, 16) = 0,1650 | P=0,6900 |
| | Genotype | 0.00001058 | 1 | 0.00001058 | F (1, 16) = 0,6561 | P=0,4298 |
| | Treatment | 0.0001935 | 1 | 0.0001935 | F (1, 16) = 12,00 | P=0,0032 |
| | Residual | 0.0002579 | 16 | 0.00001612 | | |
| **C4b** | **ANOVA table** | **SS (Type III)** | **DF** | **MS** | **F (DFn, DFd)** | **P value** |
| | Interaction | 0.00002439 | 1 | 0.00002439 | F (1, 17) = 1,605 | P=0,2223 |
| | Genotype | 2.376E-08 | 1 | 2.376E-08 | F (1, 17) = 0,00156 | P=0,9689 |
| | Treatment | 0.00001176 | 1 | 0.00001176 | F (1, 17) = 0,7742 | P=0,3912 |
| | Residual | 0.0002583 | 17 | 0.00001519 | | |
| **PUMA** | **ANOVA table** | **SS (Type III)** | **DF** | **MS** | **F (DFn, DFd)** | **P value** |
| | Interaction | 4.92E-07 | 1 | 0.000000492 | F (1, 17) = 0,9161 | P=0,3519 |
| | Genotype | 6.423E-07 | 1 | 6.423E-07 | F (1, 17) = 1,196 | P=0,2894 |
| | Treatment | 1.959E-06 | 1 | 0.000001959 | F (1, 17) = 3,647 | P=0,0732 |
| | Residual | 0.00000913 | 17 | 5.371E-07 | | |
| **TREM2** | **ANOVA table** | **SS (Type III)** | **DF** | **MS** | **F (DFn, DFd)** | **P value** |
| | Interaction | 2.897E-07 | 1 | 2.897E-07 | F (1, 15) = 1,303 | P=0,2716 |
| | Genotype | 1.314E-07 | 1 | 1.314E-07 | F (1, 15) = 0,5907 | P=0,4541 |
| | Treatment | 1.109E-06 | 1 | 0.000001109 | F (1, 15) = 4,988 | P=0,0412 |
| | Residual | 3.336E-06 | 15 | 2.224E-07 | | |

### Figure 5B Striatum

| TNF-a | ANOVA table | SS (Type III) | DF | MS | F (DFn | P value |
|---|---|---|---|---|---|---|
| | Interaction | 3.535E-11 | 1 | 0 | F (1, 1 | P=0,5490 |
| | Genotype | 2.336E-10 | 1 | 0 | F (1, 1 | P=0,1350 |
| | Treatment | 1.319E-10 | 1 | 0 | F (1, 1 | P=0,2541 |
| | Residual | 1.509E-09 | 16 | 0 | | |
| IFN-b | ANOVA table | SS (Type III) | DF | MS | F (DFn | P value |
| | Interaction | 4.93E-09 | 1 | 0 | F (1, 1 | P=0,0513 |
| | Genotype | 1.532E-12 | 1 | 0 | F (1, 1 | P=0,9708 |
| | Treatment | 7.58E-10 | 1 | 0 | F (1, 1 | P=0,4210 |
| | Residual | 1.778E-08 | 16 | 0 | | |
| IL1-b | ANOVA table | SS (Type III) | DF | MS | F (DFn | P value |
| | Interaction | 1.283E-09 | 1 | 0 | F (1, 1 | P=0,0994 |
| | Genotype | 2.544E-09 | 1 | 0 | F (1, 1 | P=0,0252 |
| | Treatment | 5.573E-09 | 1 | 0 | F (1, 1 | P=0,0021 |
| | Residual | 7.18E-09 | 17 | 0 | | |
| IL-6 | ANOVA table | SS (Type III) | DF | MS | F (DFn | P value |
| | Interaction | 7.412E-10 | 1 | 0 | F (1, 1 | P=0,0835 |
| | Genotype | 3.249E-10 | 1 | 0 | F (1, 1 | P=0,2402 |
| | Treatment | 2.335E-10 | 1 | 0 | F (1, 1 | P=0,3167 |
| | Residual | 3.728E-09 | 17 | 0 | | |
| CXCL10 | ANOVA table | SS (Type III) | DF | MS | F (DFn | P value |
| | Interaction | 4.19E-08 | 1 | 0 | F (1, 1 | P=0,5118 |
| | Genotype | 8.986E-07 | 1 | 0 | F (1, 1 | P=0,0065 |
| | Treatment | 3.262E-07 | 1 | 0 | F (1, 1 | P=0,0789 |
| | Residual | 0.000001587 | 17 | 0 | | |
| Mx1 | ANOVA table | SS (Type III) | DF | MS | F (DFn | P value |
| | Interaction | 2.452E-10 | 1 | 0 | F (1, 1 | P=0,5431 |
| | Genotype | 6.602E-10 | 1 | 0 | F (1, 1 | P=0,3228 |
| | Treatment | 3.83E-09 | 1 | 0 | F (1, 1 | P=0,0253 |
| | Residual | 1.082E-08 | 17 | 0 | | |
| Viperin | ANOVA table | SS (Type III) | DF | MS | F (DFn | P value |
| | Interaction | 9.011E-12 | 1 | 0 | F (1, 1 | P=0,9786 |
| | Genotype | 5.59E-08 | 1 | 0 | F (1, 1 | P=0,0471 |
| | Treatment | 2.689E-11 | 1 | 0 | F (1, 1 | P=0,9631 |
| | Residual | 2.074E-07 | 17 | 0 | | |
| CCL2 | ANOVA table | SS (Type III) | DF | MS | F (DFn | P value |
| | Interaction | 7.652E-11 | 1 | 0 | F (1, 1 | P=0,7321 |
| | Genotype | 4.851E-09 | 1 | 0 | F (1, 1 | P=0,0131 |
| | Treatment | 1.422E-09 | 1 | 0 | F (1, 1 | P=0,1519 |
| | Residual | 1.074E-08 | 17 | 0 | | |
| TLR4 | ANOVA table | SS (Type III) | DF | MS | F (DFn | P value |
| | Interaction | 1.778E-10 | 1 | 0 | F (1, 1 | P=0,3747 |
| | Genotype | 5.75E-10 | 1 | 0 | F (1, 1 | P=0,1195 |
| | Treamtent | 3.377E-10 | 1 | 0 | F (1, 1 | P=0,2261 |
| | Residual | 3.638E-09 | 17 | 0 | | |
| TLR2 | ANOVA table | SS (Type III) | DF | MS | F (DFn | P value |
| | Interaction | 2.003E-08 | 1 | 0 | F (1, 1 | P=0,0735 |
| | Row Factor | 1.837E-08 | 1 | 0 | F (1, 1 | P=0,0853 |
| | Column Factor | 1.494E-07 | 1 | 0 | F (1, 1 | P<0,0001 |
| | Residual | 9.361E-08 | 17 | 0 | | |
| C1q | ANOVA table | SS (Type III) | DF | MS | F (DFn | P value |
| | Interaction | 0.00001179 | 1 | 0 | F (1, 1 | P=0,7155 |
| | Genotype | 0.0003339 | 1 | 0 | F (1, 1 | P=0,0651 |
| | Treatment | 0.001095 | 1 | 0 | F (1, 1 | P=0,0024 |
| | Residual | 0.00146 | 17 | 0 | | |
| C4b | ANOVA table | SS (Type III) | DF | MS | F (DFn | P value |
| | Interaction | 0.0001736 | 1 | 0 | F (1, 1 | P=0,3216 |
| | Genotype | 0.0005695 | 1 | 0 | F (1, 1 | P=0,0819 |
| | Treatment | 0.002379 | 1 | 0 | F (1, 1 | P=0,0015 |
| | Residual | 0.002831 | 17 | 0 | | |
| PUMA | ANOVA table | SS (Type III) | DF | MS | F (DFn | P value |
| | Interaction | 5.224E-07 | 1 | 0 | F (1, 1 | P=0,3134 |
| | Genotype | 0.0000011 | 1 | 0 | F (1, 1 | P=0,1500 |
| | Treatment | 0.000002578 | 1 | 0 | F (1, 1 | P=0,0339 |
| | Residual | 0.000008229 | 17 | 0 | | |
| TREM2 | ANOVA table | SS (Type III) | DF | MS | F (DFn | P value |
| | Interaction | 0.000002837 | 1 | 0 | F (1, 1 | P=0,2837 |
| | Genotype | 0.00001182 | 1 | 0 | F (1, 1 | P=0,0373 |
| | Treatment | 0.00001802 | 1 | 0 | F (1, 1 | P=0,0126 |
| | Residual | 0.00003936 | 17 | 0 | | |

#### Figure 5C Ventral midbrain

| | ANOVA table | SS (Type III) | DF | MS | F (DFn, DFd) | P value |
|---|---|---|---|---|---|---|
| **TNF-a** | **ANOVA table** | **SS (Type III)** | **DF** | **MS** | **F (DFn, DFd)** | **P value** |
| | Interaction | 3.919E-10 | 1 | 3.919E-10 | F (1, 15) = 1,170 | P=0,2965 |
| | Genotype | 1.312E-10 | 1 | 1.312E-10 | F (1, 15) = 0,3918 | P=0,5408 |
| | Treatment | 2.939E-10 | 1 | 2.939E-10 | F (1, 15) = 0,8773 | P=0,3638 |
| | Residual | 5.024E-09 | 15 | 3.349E-10 | | |
| **IFN-b** | **ANOVA table** | **SS (Type III)** | **DF** | **MS** | **F (DFn, DFd)** | **P value** |
| | Interaction | 2.736E-09 | 1 | 2.736E-09 | F (1, 13) = 5,395 | P=0,0371 |
| | Genotype | 3.444E-11 | 1 | 3.444E-11 | F (1, 13) = 0,06791 | P=0,7985 |
| | Treatment | 2.249E-12 | 1 | 2.249E-12 | F (1, 13) = 0,004435 | P=0,9479 |
| | Residual | 6.593E-09 | 13 | 5.072E-10 | | |
| **IL1-b** | **ANOVA table** | **SS (Type III)** | **DF** | **MS** | **F (DFn, DFd)** | **P value** |
| | Interaction | 1.599E-09 | 1 | 1.599E-09 | F (1, 18) = 1,764 | P=0,2007 |
| | Genotype | 1.532E-09 | 1 | 1.532E-09 | F (1, 18) = 1,691 | P=0,2099 |
| | Treatment | 1.187E-09 | 1 | 1.187E-09 | F (1, 18) = 1,309 | P=0,2675 |
| | Residual | 1.631E-08 | 18 | 9.062E-10 | | |
| **IL-6** | **ANOVA table** | **SS (Type III)** | **DF** | **MS** | **F (DFn, DFd)** | **P value** |
| | Interaction | 2.878E-10 | 1 | 2.878E-10 | F (1, 18) = 1,272 | P=0,2742 |
| | Genotype | 1.089E-09 | 1 | 1.089E-09 | F (1, 18) = 4,814 | P=0,0416 |
| | Treatment | 4.105E-10 | 1 | 4.105E-10 | F (1, 18) = 1,814 | P=0,1947 |
| | Residual | 4.073E-09 | 18 | 2.263E-10 | | |
| **CXCL10** | **ANOVA table** | **SS (Type III)** | **DF** | **MS** | **F (DFn, DFd)** | **P value** |
| | Interaction | 2.242E-08 | 1 | 2.242E-08 | F (1, 17) = 1,018 | P=0,3272 |
| | Genotype | 1.643E-07 | 1 | 1.643E-07 | F (1, 17) = 7,459 | P=0,0142 |
| | Treatment | 1.913E-09 | 1 | 1.913E-09 | F (1, 17) = 0,08687 | P=0,7718 |
| | Residual | 3.744E-07 | 17 | 2.202E-08 | | |
| **Mx1** | **ANOVA table** | **SS (Type III)** | **DF** | **MS** | **F (DFn, DFd)** | **P value** |
| | Interaction | 4.104E-10 | 1 | 4.104E-10 | F (1, 18) = 0,6564 | P=0,4284 |
| | Genotype | 7.731E-15 | 1 | 7.731E-15 | F (1, 18) = 1,236e-005 | P=0,9972 |
| | Treatment | 2.013E-09 | 1 | 2.013E-09 | F (1, 18) = 3,220 | P=0,0895 |
| | Residual | 1.125E-08 | 18 | 6.252E-10 | | |
| **Viperin** | **ANOVA table** | **SS (Type III)** | **DF** | **MS** | **F (DFn, DFd)** | **P value** |
| | Interaction | 2.123E-08 | 1 | 2.123E-08 | F (1, 18) = 3,096 | P=0,0955 |
| | Genotype | 7.752E-08 | 1 | 7.752E-08 | F (1, 18) = 11,31 | P=0,0035 |
| | Treatment | 2.504E-08 | 1 | 2.504E-08 | F (1, 18) = 3,652 | P=0,0721 |
| | Residual | 1.234E-07 | 18 | 6.856E-09 | | |
| **CCL2** | **ANOVA table** | **SS (Type III)** | **DF** | **MS** | **F (DFn, DFd)** | **P value** |
| | Interaction | 2.639E-09 | 1 | 2.639E-09 | F (1, 18) = 1,167 | P=0,2943 |
| | Genotype | 4.603E-10 | 1 | 4.603E-10 | F (1, 18) = 0,2035 | P=0,6573 |
| | Treatment | 5.832E-11 | 1 | 5.832E-11 | F (1, 18) = 0,02579 | P=0,8742 |
| | Residual | 4.071E-08 | 18 | 2.262E-09 | | |
| **TLR4** | **ANOVA table** | **SS (Type III)** | **DF** | **MS** | **F (DFn, DFd)** | **P value** |
| | Interaction | 5.092E-10 | 1 | 5.092E-10 | F (1, 17) = 1,264 | P=0,2765 |
| | Genotype | 7.406E-10 | 1 | 7.406E-10 | F (1, 17) = 1,839 | P=0,1928 |
| | Treamtent | 4.062E-10 | 1 | 4.062E-10 | F (1, 17) = 1,009 | P=0,3293 |
| | Residual | 6.846E-09 | 17 | 4.027E-10 | | |
| **TLR2** | **ANOVA table** | **SS (Type III)** | **DF** | **MS** | **F (DFn, DFd)** | **P value** |
| | Interaction | 3.79E-10 | 1 | 3.79E-10 | F (1, 17) = 0,2847 | P=0,6005 |
| | Row Factor | 5.133E-09 | 1 | 5.133E-09 | F (1, 17) = 3,856 | P=0,0661 |
| | Column Factor | 4.174E-10 | 1 | 4.174E-10 | F (1, 17) = 0,3136 | P=0,5828 |
| | Residual | 2.263E-08 | 17 | 1.331E-09 | | |
| **C1q** | **ANOVA table** | **SS (Type III)** | **DF** | **MS** | **F (DFn, DFd)** | **P value** |
| | Interaction | 0.00005987 | 1 | 0.00005987 | F (1, 17) = 5,770 | P=0,0280 |
| | Genotype | 0.000202 | 1 | 0.000202 | F (1, 17) = 19,47 | P=0,0004 |
| | Treatment | 0.00007299 | 1 | 0.00007299 | F (1, 17) = 7,034 | P=0,0168 |
| | Residual | 0.0001764 | 17 | 0.00001038 | | |
| **C4b** | **ANOVA table** | **SS (Type III)** | **DF** | **MS** | **F (DFn, DFd)** | **P value** |
| | Interaction | 0.00005984 | 1 | 0.00005984 | F (1, 17) = 2,139 | P=0,1619 |
| | Genotype | 0.000126 | 1 | 0.000126 | F (1, 17) = 4,504 | P=0,0488 |
| | Treatment | 0.00007249 | 1 | 0.00007249 | F (1, 17) = 2,591 | P=0,1259 |
| | Residual | 0.0004756 | 17 | 0.00002798 | | |
| **PUMA** | **ANOVA table** | **SS (Type III)** | **DF** | **MS** | **F (DFn, DFd)** | **P value** |
| | Interaction | 0.000009358 | 1 | 0.000009358 | F (1, 18) = 3,582 | P=0,0746 |
| | Genotype | 0.000005413 | 1 | 0.000005413 | F (1, 18) = 2,072 | P=0,1672 |
| | Treatment | 0.000004897 | 1 | 0.000004897 | F (1, 18) = 1,874 | P=0,1878 |
| | Residual | 0.00004702 | 18 | 0.000002612 | | |
| **TREM2** | **ANOVA table** | **SS (Type III)** | **DF** | **MS** | **F (DFn, DFd)** | **P value** |
| | Interaction | 0.000001649 | 1 | 0.000001649 | F (1, 17) = 2,527 | P=0,1303 |
| | Genotype | 0.000003665 | 1 | 0.000003665 | F (1, 17) = 5,616 | P=0,0299 |
| | Treatment | 0.000002875 | 1 | 0.000002875 | F (1, 17) = 4,405 | P=0,0511 |
| | Residual | 0.00001109 | 17 | 6.526E-07 | | |

#### Figure 6B

##### 1 month

| MJF14 %area Amygdala | ANOVA table | SS (Type III) | DF | MS | F (DFn, DFd) | P value |
|---|---|---|---|---|---|---|
| | Interaction | 0,02261 | 1 | 0,023 | F (1, 33) = 0,9474 | P=0,3375 |
| | Genotype | 0,02864 | 1 | 0,029 | F (1, 33) = 1,200 | P=0,2812 |
| | Treatment | 0,4234 | 1 | 0,423 | F (1, 33) = 17,74 | P=0,0002 |
| | Residual | 0,7875 | 33 | 0,024 | | |

##### 6 months

| MJF14 %area Amygdala | ANOVA table | SS (Type III) | DF | MS | F (DFn, DFd) | P value |
|---|---|---|---|---|---|---|
| | Interaction | 0,00002504 | 1 | 0,00002504 | F (1, 21) = 0,00021 | P=0,9885 |
| | Genotype | 8,556E-06 | 1 | 0,000008556 | F (1, 21) = 7,213e- | P=0,9933 |
| | Treatment | 1,393 | 1 | 1,393 | F (1, 21) = 11,74 | P=0,0025 |
| | Residual | 2,491 | 21 | 0,1186 | | |

#### Figure 6C

##### 1 month

| MJF14 %area SN | ANOVA table | SS (Type III) | DF | MS | F (DFn, DFd) | P value |
|---|---|---|---|---|---|---|
| | Interaction | 0,0003403 | 1 | 0,0003403 | F (1, 21) = 0,04094 | P=0,8416 |
| | Genotype | 0,0004057 | 1 | 0,0004057 | F (1, 21) = 0,04880 | P=0,8273 |
| | Treatment | 0,05297 | 1 | 0,05297 | F (1, 21) = 6,371 | P=0,0197 |
| | Residual | 0,1746 | 21 | 0,008313 | | |

##### 6 months

| MJF14 %area SN | ANOVA table | SS (Type III) | DF | MS | F (DFn, DFd) | P value |
|---|---|---|---|---|---|---|
| | Interaction | 0,0006161 | 1 | 6E-04 | F (1, 33) = 1,030 | P=0,3176 |
| | Genotype | 0,00005385 | 1 | 5E-05 | F (1, 33) = 0,09000 | P=0,7661 |
| | Treatment | 0,0138 | 1 | 0,014 | F (1, 33) = 23,07 | P<0,0001 |
| | Residual | 0,01974 | 33 | 6E-04 | | |

#### Figure 6D

##### 1 month

| p62 %area Amygdala | ANOVA table | SS (Type III) | DF | MS | F (DFn, DFd) | P value |
|---|---|---|---|---|---|---|
| | Interaction | 0,000116 | 1 | 0,000116 | F (1, 19) = 0,00162 | P=0,9683 |
| | Genotype | 0,0006095 | 1 | 0,0006095 | F (1, 19) = 0,00850 | P=0,9275 |
| | Treatment | 0,4415 | 1 | 0,4415 | F (1, 19) = 6,164 | P=0,0225 |
| | Residual | 1,361 | 19 | 0,07164 | | |

##### 6 months

| p62 %area Amygdala | ANOVA table | SS (Type III) | DF | MS | F (DFn, DFd) | P value |
|---|---|---|---|---|---|---|
| | Interaction | 0,00231 | 1 | 0,002 | F (1, 28) = 0,3001 | P=0,5882 |
| | Genotype | 0,007669 | 1 | 0,008 | F (1, 28) = 0,9961 | P=0,3268 |
| | Treatment | 0,2225 | 1 | 0,223 | F (1, 28) = 28,90 | P<0,0001 |
| | Residual | 0,2156 | 28 | 0,008 | | |

#### Figure 6E

##### 1 month

| p62 %area SN | ANOVA table | SS (Type III) | DF | MS | F (DFn, DFd) | P value |
|---|---|---|---|---|---|---|
| | Interaction | 0,00000103 | 1 | 0,00000103 | F (1, 21) = 0,00456 | P=0,9468 |
| | Genotype | 0,0000751 | 1 | 0,0000751 | F (1, 21) = 0,3327 | P=0,5702 |
| | Treatment | 0,005778 | 1 | 0,005778 | F (1, 21) = 25,60 | P<0,0001 |
| | Residual | 0,00474 | 21 | 0,0002257 | | |

##### 6 months

| p62 %area SN | ANOVA table | SS (Type III) | DF | MS | F (DFn, DFd) | P value |
|---|---|---|---|---|---|---|
| | Interaction | 0,00001757 | 1 | 2E-05 | F (1, 33) = 0,03445 | P=0,8539 |
| | Genotypes | 4,516E-06 | 1 | 5E-06 | F (1, 33) = 0,008853 | P=0,9256 |
| | Treatment | 0,01277 | 1 | 0,013 | F (1, 33) = 25,04 | P<0,0001 |
| | Residual | 0,01683 | 33 | 5E-04 | | |

#### Figure 6F

| Pser129 %area Amygdala | ANOVA table | SS (Type III) | DF | MS | F (DFn, DFd) | P value |
|---|---|---|---|---|---|---|
| | Interaction | 0,4314 | 2 | 0,216 | F (2, 38) = 0,8626 | P=0,4301 |
| | Treatment | 5,86 | 2 | 2,93 | F (2, 38) = 11,72 | P=0,0001 |
| | Genotype | 0,2651 | 1 | 0,265 | F (1, 38) = 1,060 | P=0,3097 |
| | Residual | 9,502 | 38 | 0,25 | | |

#### Fig 6G

| Pser129 %Area SN | ANOVA table | SS (Type III) | DF | MS | F (DFn, DFd) | P value |
|---|---|---|---|---|---|---|
| | Interaction | 0.003383 | 2 | 0.001692 | F (2, 36) = 0,3490 | P=0,7078 |
| | Treatment | 0.3849 | 2 | 0.1924 | F (2, 36) = 39,70 | P<0,0001 |
| | Genotype | 0.005369 | 1 | 0.005369 | F (1, 36) = 1,108 | P=0,2996 |
| | Residual | 0.1745 | 36 | 0.004847 | | |

#### Figure 7B

| TH OD 1 month (%contralateral) | ANOVA table | SS (Type III) | DF | MS | F (DFn, DFd) | P value |
|---|---|---|---|---|---|---|
| | Interaction | 152,4 | 2 | 76,19 | F (2, 28) = 1,541 | P=0,2317 |
| | Treatment | 1472 | 2 | 736,2 | F (2, 28) = 14,90 | P<0,0001 |
| | Genotype | 94,08 | 1 | 94,08 | F (1, 28) = 1,903 | P=0,1786 |
| | Residual | 1384 | 28 | 49,43 | | |

#### Figure 7C

| TH OD 6 months (%contralateral) | ANOVA table | SS (Type III) | DF | MS | F (DFn, DFd) | P value |
|---|---|---|---|---|---|---|
| | Interaction | 57,16 | 2 | 28,58 | F (2, 42) = 0,4553 | P=0,6374 |
| | Treatment | 7695 | 2 | 3848 | F (2, 42) = 61,29 | P<0,0001 |
| | Genotype | 0,6648 | 1 | 0,6648 | F (1, 42) = 0,01059 | P=0,9185 |
| | Residual | 2637 | 42 | 62,78 | | |

#### Figure 8B

| TH %contralateral 1 month | ANOVA table | SS (Type III) | DF | MS | F (DFn, DFd) | P value |
|---|---|---|---|---|---|---|
| | Interaction | 127,8 | 2 | 63,89 | F (2, 28) = 0,8275 | P=0,4475 |
| | Genotype | 59,05 | 1 | 59,05 | F (1, 28) = 0,7649 | P=0,3892 |
| | Treatment | 1413 | 2 | 706,6 | F (2, 28) = 9,152 | P=0,0009 |
| | Residual | 2162 | 28 | 77,2 | | |

#### Figure 8C

| TH %contralateral 6 months | ANOVA table | SS (Type III) | DF | MS | F (DFn, DFd) | P value |
|---|---|---|---|---|---|---|
| | Interaction | 19,25 | 2 | 9,623 | F (2, 43) = 0,08459 | P=0,9190 |
| | Genotype | 111,5 | 1 | 111,5 | F (1, 43) = 0,9799 | P=0,3278 |
| | Treatment | 16624 | 2 | 8312 | F (2, 43) = 73,07 | P<0,0001 |
| | Residual | 4892 | 43 | 113,8 | | |

#### Supplementary Figure 1A

| Challening beam Total Errors/steps | ANOVA table | SS (Type III) | DF | MS | F (DFn, DFd) | P value |
|---|---|---|---|---|---|---|
| | Interaction | 0,001788 | 2 | 0,0008942 | F (2, 60) = 0,02486 | P=0,9755 |
| | Genotype | 0,04319 | 1 | 0,04319 | F (1, 60) = 1,201 | P=0,2776 |
| | Treatment | 0,4131 | 2 | 0,2065 | F (2, 60) = 5,742 | P=0,0052 |
| | Residual | 2,158 | 60 | 0,03597 | | |
| **Challening beam Total steps/Sec** | **ANOVA table** | **SS (Type III)** | **DF** | **MS** | **F (DFn, DFd)** | **P value** |
| | Interaction | 2,471 | 2 | 1,236 | F (2, 61) = 0,5433 | P=0,5836 |
| | Genotype | 20,16 | 1 | 20,16 | F (1, 61) = 8,866 | P=0,0042 |
| | Treatment | 64,69 | 2 | 32,35 | F (2, 61) = 14,22 | P<0,0001 |
| | Residual | 138,7 | 61 | 2,274 | | |

#### Supplementary figure 1B

| Challening beam Total Errors/steps | ANOVA table | SS (Type III) | DF | MS | F (DFn, DFd) | P value |
|---|---|---|---|---|---|---|
| | Interaction | 0,3521 | 2 | 0,176 | F (2, 46) = 3,123 | P=0,0535 |
| | Genotype | 0,2666 | 1 | 0,267 | F (1, 46) = 4,730 | P=0,0348 |
| | Treatment | 0,628 | 2 | 0,314 | F (2, 46) = 5,571 | P=0,0068 |
| | Residual | 2,593 | 46 | 0,056 | | |
| **Challening beam Total steps/Sec** | **ANOVA table** | **SS (Type III)** | **DF** | **MS** | **F (DFn, DFd)** | **P value** |
| | Interaction | 10,95 | 2 | 5,477 | F (2, 46) = 2,710 | P=0,0771 |
| | Genotype | 4,413 | 1 | 4,413 | F (1, 46) = 2,184 | P=0,1463 |
| | Treatment | 58,91 | 2 | 29,45 | F (2, 46) = 14,58 | P<0,0001 |
| | Residual | 92,95 | 46 | 2,021 | | |

#### Supplementary figure 2A

| Rearings per minute | ANOVA table | SS (Type III) | DF | MS | F (DFn, DFd) | P value |
|---|---|---|---|---|---|---|
| | Interaction | 26.05 | 2 | 13.03 | F (2, 45) = 1,659 | P=0,2017 |
| | Genotype | 0.6933 | 1 | 0.693 | F (1, 45) = 0,08829 | P=0,7677 |
| | Treatment | 65.94 | 2 | 32.97 | F (2, 45) = 4,199 | P=0,0213 |
| | Residual | 353.3 | 45 | 7.852 | | |

#### Supplementary figure 2B

| Grooming (time) | ANOVA table | SS (Type III) | DF | MS | F (DFn, DFd) | P value |
|---|---|---|---|---|---|---|
| | Interaction | 6.478 | 2 | 3.239 | F (2, 45) = 0,3492 | P=0,7071 |
| | Genotype | 23.88 | 1 | 23.88 | F (1, 45) = 2,575 | P=0,1155 |
| | Treatment | 43.29 | 2 | 21.65 | F (2, 45) = 2,334 | P=0,1085 |
| | Residual | 417.3 | 45 | 9.274 | | |

#### Supplementary figure 2C

| Forelimbs: Left and Right | ANOVA table | SS | DF | MS | F (DFn, DFd) | P value | Repeate measure |
|---|---|---|---|---|---|---|---|
| | Treatment | 36083 | 2 | 18041 | F (2, 46) = 10,87 | P=0,0001 | |
| | Genotype | 5037 | 1 | 5037 | F (1, 46) = 3,036 | P=0,0881 | |
| | Left/Right | 208.5 | 1 | 208.5 | F (1, 46) = 1,915 | P=0,1730 | |
| | Treatment x Genotype | 4477 | 2 | 2239 | F (2, 46) = 1,349 | P=0,2695 | |
| | Treatment x Left/Right | 708.1 | 2 | 354.1 | F (2, 46) = 3,252 | P=0,0477 | |
| | Genotype x Left/Right | 170.9 | 1 | 170.9 | F (1, 46) = 1,570 | P=0,2165 | |
| | Treatment x Genotype x Left/Right | 214.4 | 2 | 107.2 | F (2, 46) = 0,9848 | P=0,3812 | |
| | Subject | 76327 | 46 | 1659 | | | |
| | Residual | 5008 | 46 | 108.9 | | | |

#### Supplementary figure 2D

| Hindlimbs: Left and Right | ANOVA table | SS | DF | MS | F (DFn, DFd) | P value | Repeate measure |
|---|---|---|---|---|---|---|---|
| | Treatment | 33953 | 2 | 16977 | F (2, 46) = 6,622 | P=0,0030 | |
| | Genotype | 4295 | 1 | 4295 | F (1, 46) = 1,675 | P=0,2020 | |
| | Left/right | 1802 | 1 | 1802 | F (1, 46) = 17,44 | P=0,0001 | |
| | Treatment x Genotype | 3621 | 2 | 1811 | F (2, 46) = 0,7062 | P=0,4988 | |
| | Treatment x Left/right | 365.6 | 2 | 182.8 | F (2, 46) = 1,769 | P=0,1819 | |
| | Genotype x Left/right | 48.83 | 1 | 48.83 | F (1, 46) = 0,4725 | P=0,4953 | |
| | Treatment x Genotype x Left/right | 531.7 | 2 | 265.9 | F (2, 46) = 2,573 | P=0,0873 | |
| | Subject | 117933 | 46 | 2564 | | | |
| | Residual | 4754 | 46 | 103.3 | | | |

#### Supplementary figure 3A

| Type A (repeated measure) | ANOVA table | SS (Type III) | DF | MS | F (DFn, DFd) | P value |
|---|---|---|---|---|---|---|
| | Treatment | 4859071 | 2 | 2429536 | F (2, 27) = 5,614 | P=0,0091 |
| | Genotype | 357977 | 1 | 357977 | F (1, 27) = 0,8271 | P=0,3712 |
| | Hemisphere | 80154 | 1 | 80154 | F (1, 27) = 0,3182 | P=0,5774 |
| | Treatment x Genotype | 2941611 | 2 | 1470806 | F (2, 27) = 3,398 | P=0,0483 |
| | Treatment x Hemisphere | 209835 | 2 | 104918 | F (2, 27) = 0,4165 | P=0,6635 |
| | Genotype x Hemisphere | 778317 | 1 | 778317 | F (1, 27) = 3,090 | P=0,0901 |
| | Treatment x Genotype x Hemisphere | 71123 | 2 | 35561 | F (2, 27) = 0,1412 | P=0,8690 |
| | animal | 11685605 | 27 | 432800 | | |
| | Residual | 6801887 | 27 | 251922 | | |
| **Type B (repeated measure)** | **ANOVA table** | **SS (Type III)** | **DF** | **MS** | **F (DFn, DFd)** | **P value** |
| | Treatment | 2471248 | 2 | 1235624 | F (2, 27) = 1,750 | P=0,1929 |
| | Genotype | 12641 | 1 | 12641 | F (1, 27) = 0,01791 | P=0,8945 |
| | Hemisphere | 465116 | 1 | 465116 | F (1, 27) = 1,648 | P=0,2101 |
| | Treatment x Genotype | 450410 | 2 | 225205 | F (2, 27) = 0,3190 | P=0,7296 |
| | Treatment x Hemisphere | 42136 | 2 | 21068 | F (2, 27) = 0,07467 | P=0,9282 |
| | Genotype x Hemisphere | 5222 | 1 | 5222 | F (1, 27) = 0,01851 | P=0,8928 |
| | Treatment x Genotype x Hemisphere | 91863 | 2 | 45931 | F (2, 27) = 0,1628 | P=0,8506 |
| | animal | 19060165 | 27 | 705932 | | |
| | Residual | 7618084 | 27 | 282151 | | |
| **Type C (repeated measure)** | **ANOVA table** | **SS (Type III)** | **DF** | **MS** | **F (DFn, DFd)** | **P value** |
| | ANOVA table | SS | DF | MS | F (DFn, DFd) | P value |
| | Treatment | 2101033 | 2 | 1050516 | F (2, 27) = 8,157 | P=0,0017 |
| | Genotype | 13191 | 1 | 13191 | F (1, 27) = 0,1024 | P=0,7514 |
| | Hemisphere | 177411 | 1 | 177411 | F (1, 27) = 2,545 | P=0,1223 |
| | Treatment x Genotype | 38081 | 2 | 19040 | F (2, 27) = 0,1479 | P=0,8633 |
| | Treatment x Hemisphere | 140945 | 2 | 70473 | F (2, 27) = 1,011 | P=0,3772 |
| | Genotype x Hemisphere | 10154 | 1 | 10154 | F (1, 27) = 0,1457 | P=0,7057 |
| | Treatment x Genotype x Hemisphere | 64586 | 2 | 32293 | F (2, 27) = 0,4633 | P=0,6341 |
| | animal | 3477077 | 27 | 128781 | | |
| | Residual | 1881996 | 27 | 69704 | | |

#### Supplementary figure 3B

| Type A (repeated measure) | ANOVA table | SS (Type III) | DF | MS | F (DFn, DFd) | P value |
|---|---|---|---|---|---|---|
| | treatment | 17392547 | 2 | 8696274 | F (2, 46) = 17,46 | P<0,0001 |
| | Genotype | 12474 | 1 | 12474 | F (1, 46) = 0,02505 | P=0,8749 |
| | Hemisphere | 6772484 | 1 | 6772484 | F (1, 46) = 32,17 | P<0,0001 |
| | treatment x Genotype | 1402247 | 2 | 701123 | F (2, 46) = 1,408 | P=0,2550 |
| | treatment x Hemisphere | 5128121 | 2 | 2564060 | F (2, 46) = 12,18 | P<0,0001 |
| | Genotype x Hemisphere | 110709 | 1 | 110709 | F (1, 46) = 0,5259 | P=0,4720 |
| | treatment x Genotype x Hemisphere | 99846 | 2 | 49923 | F (2, 46) = 0,2371 | P=0,7898 |
| | animal | 22909356 | 46 | 498029 | | |
| | Residual | 9684041 | 46 | 210523 | | |
| **Type B (repeated measure)** | **ANOVA table** | **SS (Type III)** | **DF** | **MS** | **F (DFn, DFd)** | **P value** |
| | Row factor | 2544736 | 2 | 1272368 | F (2, 46) = 1,737 | P=0,1875 |
| | (AB vs CD) | 1628150 | 1 | 1628150 | F (1, 46) = 2,222 | P=0,1429 |
| | (AC vs BD) | 9536184 | 1 | 9536184 | F (1, 46) = 31,57 | P<0,0001 |
| | Row factor x (AB vs CD) | 5387835 | 2 | 2693917 | F (2, 46) = 3,677 | P=0,0330 |
| | Row factor x (AC vs BD) | 2558867 | 2 | 1279433 | F (2, 46) = 4,236 | P=0,0205 |
| | (AB vs CD) x (AC vs BD) | 33488 | 1 | 33488 | F (1, 46) = 0,1109 | P=0,7407 |
| | Row factor x (AB vs CD) x (AC vs BI | 230764 | 2 | 115382 | F (2, 46) = 0,3820 | P=0,6846 |
| | Subject | 33704663 | 46 | 732710 | | |
| | Residual | 13894306 | 46 | 302050 | | |
| **Type C (repeated measure)** | **ANOVA table** | **SS (Type III)** | **DF** | **MS** | **F (DFn, DFd)** | **P value** |
| | treatment | 65183663 | 2 | 32591831 | F (2, 46) = 61,37 | P<0,0001 |
| | Genotype | 221,5 | 1 | 221,5 | F (1, 46) = 0,0004172 | P=0,9838 |
| | Hemisphere | 16459298 | 1 | 16459298 | F (1, 46) = 65,30 | P<0,0001 |
| | treatment x Genotype | 89282 | 2 | 44641 | F (2, 46) = 0,08406 | P=0,9195 |
| | treatment x Hemisphere | 8318625 | 2 | 4159313 | F (2, 46) = 16,50 | P<0,0001 |
| | Genotype x Hemisphere | 63736 | 1 | 63736 | F (1, 46) = 0,2529 | P=0,6175 |
| | treatment x Genotype x Hemisphere | 35940 | 2 | 17970 | F (2, 46) = 0,07129 | P=0,9313 |
| | animal | 24427658 | 46 | 531036 | | |
| | Residual | 11595197 | 46 | 252070 | | |

#### Supplementary figure 6B

| Pser129 %area frontal cortex | ANOVA table | SS (Type III) | DF | MS | F (DFn, DFd) | P value |
|---|---|---|---|---|---|---|
| | Interaction | 0,08823 | 2 | 0,044 | F (2, 39) = 1,190 | P=0,3150 |
| | Treatment | 1,274 | 2 | 0,637 | F (2, 39) = 17,18 | P<0,0001 |
| | Genotype | 0,04288 | 1 | 0,043 | F (1, 39) = 1,157 | P=0,2887 |
| | Residual | 1,446 | 39 | 0,037 | | |

Supplementary figure 6C

| Pser129 %area Thalamus | ANOVA table | SS (Type III) | DF | MS | F (DFn, DFd) | P value |
|---|---|---|---|---|---|---|
| | Interaction | 0,002664 | 2 | 0,001 | F (2, 39) = 0,2714 | P=0,7638 |
| | Treatment | 0,2937 | 2 | 0,147 | F (2, 39) = 29,91 | P<0,0001 |
| | Genotype | 0,001963 | 1 | 0,002 | F (1, 39) = 0,3998 | P=0,5309 |
| | Residual | 0,1915 | 39 | 0,005 | | |
